# Classification of dinosaur footprints using machine learning

**DOI:** 10.1101/2024.07.15.603597

**Authors:** Michael Jones, Jens N. Lallensack, Ian Jarman, Peter Falkingham, Ivo Siekmann

**Author notes:** Corresponding author: School of Computer Science and Mathematics (CSM), Faculty of Engineering and Technology (FET), Liverpool John Moores University (LJMU), Liverpool, James Parsons Building, Byrom Way, L3 3AF, United Kingdom,. RH: Jones et al.—ML classification of footprints. SUPPLEMENTARY FILE(S)––Supplementary file(s) are available for this article for free at www.tandfonline.com/UJVP.

## Abstract

Fossilised dinosaur footprints enable us to study the behaviour of individual dinosaurs as well as interactions between dinosaurs of the same or different species. There are two principal groups of three-toed dinosaurs, ornithopods and theropods. Determining if a footprint is from an ornithopod or a theropod is a challenging problem. Based on a data set of over 300 dinosaur footprints we train several machine learning models for classifying footprints as either ornithopods or theropods. The data are provided in the form of 20 landmarks for representing each footprint which are derived from images. Variable selection using logistic forward regression demonstrates that the selected landmarks are at locations that are intuitively expected to be especially informative locations, such as the top or the bottom of a footprint. Most models show good accuracy but the recall of ornithopods, of which fewer samples were contained in the data set, was generally lower than the recall of theropods. The Multi-Layer Perceptron (MLP) stands out as the model which did best at dealing with the class imbalance. Finally, we investigate which footprints were misclassified by the majority of models. We find that some misclassified samples exhibit features that are characteristic of the other class or have a compromised shape, for example, a middle toe that points to the left or the right rather than straight ahead.

## INTRODUCTION

Ornithopods and theropods are two groups of tridactyl (three-toed) dinosaurs. Whereas the herbivorous ornithopods went extinct at the end of the Cretaceous period, the primarily carnivorous theropods evolved into modern birds. Well-known examples of theropods include *Tyrannosaurus rex* and the proto-bird *Archaeopteryx*, one of the best-known ornithopods is *Iguanodon*. Fossilised footprints are a valuable data source that can provide insight into the speed at which a dinosaur moved as well as more general traits of their behaviour. Throughout the literature they have predominantly been reported as photographic images and outline tracings, though this has changed in recent years via the aquistion of 3D data (Falkingham et al 2018). However, even when full 3D data are collected, these images are then often further simplified for statistical analysis by extracting morphological features such as lengths, widths, and angles.

Broadly, the shapes of ornithopod and theropod footprints differ in various characteristics. Ornithopod footprints are usually wider and more symmetric than theropod footprints. Also, their middle toes (digit III) are commonly shorter i.e. they do not extend as far beyond the other toes (digits II and IV) as in theropods. However, these observations are unsuitable for defining general rules for distinguishing ornithopods and theropods because the characteristics described above can be found in both groups. Moreover, as Falkingham (2014) shows, the shape of a footprint is not only determined by the anatomy of the foot but also by the properties of the substrate as well as the dynamics of the dinosaur as it left the footprint; they refer to the influence of these three dimensions as the morphospace. Thus, determining if a given footprint is from an ornithopod or a theropod remains a difficult problem.

A prime example of this difficulty lies with the large tridactyl prints of Lark Quarry, Australia (Thulborn & Wade, 1979). Originally described as theropod in origin (Thulborn & Wade, 1979; Thulborn & Wade, 1984), the tracks were later reinterpreted by Romilio and Salisbury (2011) and Romilio et al. (2013) as having been made by an Ornithopod. Key to this reinterpreation was Romilio and colleagues’ use of multivariate analysis techniques pioneered by Moratalla et al. (1988). This was later critiqued by Thulborn (2013), whose critique was again then questioned by Romilio and Salisbury (2014). Falkingham et al. (2016) subsequently demonstrated that even applying the ostensibly ‘objective’ multivariate analysis resulted in highly subjective interpretations depending on where in the 3D geometry an ‘outline’ was picked, stressing that using a single outline in this way was flawed. Falkingham ultimately determined that the tracks were more theropod-like than ornithopod-like closer to the foot-sediment interface. These same tracks were revisited in Lallensack et al.’s first application of machine learning (Lallensack et al., 2022), where interpreted track-maker varied depending on which track, and whose outline was passed to the algorithm.

The aim of our study is to address this challenge by designing classifiers based on several machine learning methods. This has the additional benefit that the decision if a footprint is classified as an ornithopod or a theropod is based on statistical analysis of the data rather than on subjective decisions.

Several representative publications classify footprints by identifying clusters in scatter plots of various metrics (Castanera et al., 2013; Demathieu, 1990; dePolo et al., 2020; Figueiredo et al., 2017; Mateus & Milàn, 2008; Piñuela et al., 2016; Romilio & Salisbury, 2011; Schulp & Al-Wosabi, 2012; Thulborn, 2013) which relies on subjective decisions such as the selection of metrics to be considered as well as the definition of the cluster boundaries. The quantitative analysis of differences between ornithopods and theropods begins with the seminal study by Moratalla et al. (1988). The authors selected a sample of 66 tridactyl footprints from the Lower Cretaceous period and originating from various geographical locations. They applied factor analysis (FA) and linear discriminant analysis (LDA) to features such as length and width of each digit with the aim to discriminate between theropods and ornithopods.

Lallensack et al. (2016) looked at three trackways from the Lower Cretaceous period found in Münchehagen Germany. The authors applied geometric morphometrics to footprint outlines and landmarks. They performed Principal Component Analysis (PCA) and found that asymmetry in the terminations of the digit impressions were a large distinguishing factor between the two groups. Lallensack (2019) extended this work by developing an algorithm that could automatically define the outline of a footprint. They tested this with a single theropod trackway from the same time period and geographical region as the trackways used by Lallensack et al. (2016). Lallensack et al. (2020) aimed to characterise the variability of footprint shapes over time and between theropods and ornithopods. The authors took 303 footprints originating from the Late Triassic period, through the Jurassic all the way to the Late Cretaceous period from 134 publications. Each footprint was described by 34 landmarks and reference points and information such as the size was also recorded. They observed that ornithopod footprints are, on average, larger and wider than theropod footprints. Moreover, the features distinguishing theropods and ornithopods change with size, and the small ornithischian footprints are most similar in shape to large theropod footprints. Ornithischian footprints increased in size over time from the early Jurassic to the late Cretaceous period indicating an increase in body size over that time period, which, interestingly, is not observed in theropod footprints.

The data by Lallensack et al. (2020) will be used in this study for developing classification algorithms that enable us to automatically distinguish between ornithopod and theropod footprints.

## MATERIALS AND METHODS

The classification methods developed in this study are based on a data set that represents the visual features of dinosaur footprints by a system of landmarks (Lallensack et al., 2020). Lallensack et al. (2020) collected images of 303 dinosaur footprints from 134 different publications and represented the outline of each dinosaur footprint by 20 landmarks, see FIGURE 1. Two of the 303 footprints were not labelled as ornithopods or theropods, so these two samples were omitted. The remaining data set consists of 301 footprints, 108 labelled ornithopods, 193 labelled theropods. As can be seen in FIGURE 1, each of the 20 landmarks is represented as a point (*x*, *y*) in a two-dimensional coordinate system. We simplify the data set by transforming each landmark (*x*, *y*) to its distance *r* from the centre (0, 0) as illustrated by the arrows in FIGURE 1:

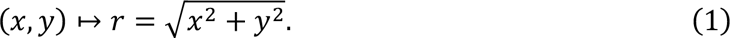

Thus, the dimensionality of the data set is reduced from 2 × 20 = 40 to 20 variables. This transformed data set, see FIGURE 2, was then used for training models for classifying footprints as theropods or ornithopods, respectively. Please note that for the purposes of this visualisation the sequence of distances from the centre plotted in FIGURE 2 was changed, starting with landmark 4 on the left of the x axis and then moving counter-clockwise along the boundary of the footprint leading to the sequence 4,7,6,5,1,8,9,20,19,18,2,15,16,17,14,13,3,10,11,12 (cf. FIGURE 1).

The mean distances of each landmark for ornithopods and theropods (FIGURE 2a) already give some insight into the differences between the two classes. When comparing the red curve representing the ornithopods and the black curve showing the theropods, landmarks 1, 5, 6, 7 – on the right of the footprint, see FIGURE 1 – and landmarks 3, 10, 11 and 12 – on the left of the footprint, see FIGURE 1 – are, on average, further away from the centre for ornithopod footprints than for theropod footprints. These observations illustrate that ornithopod footprints are usually wider than theropod footprints. Carrying out a multivariate test that extends the two-sample t-test to multiple variables as described in Härdle and Simar (2014) it can be shown that the mean distances of the landmarks are significantly different (α = 0.05) for ornithopods and theropods.

**FIGURE 1.**
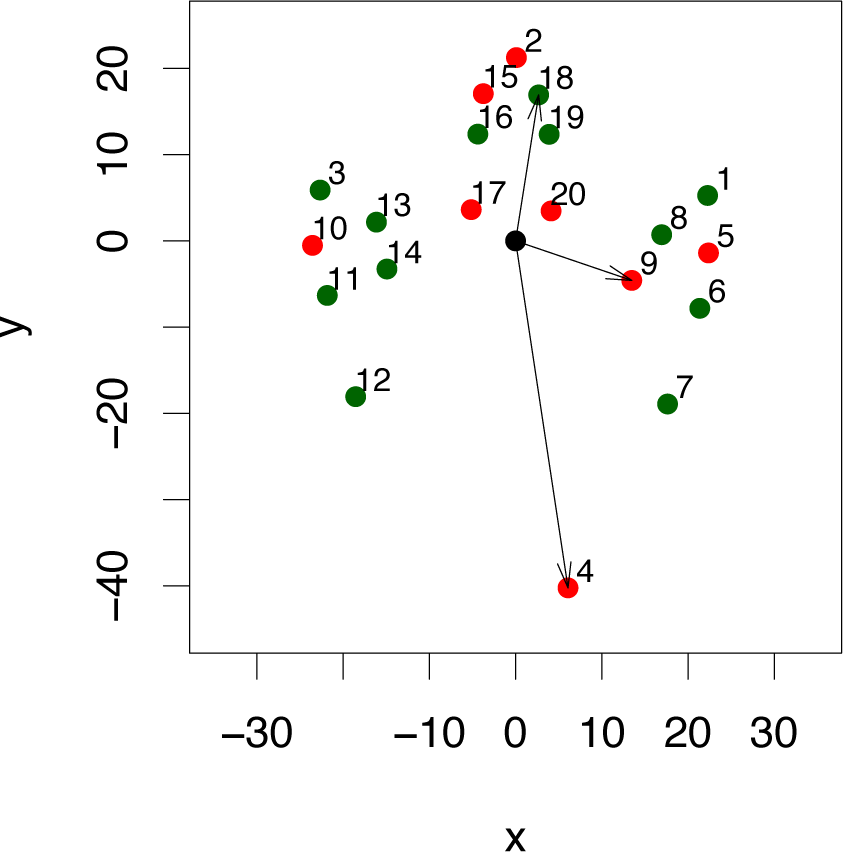
Example for a footprint from the data set by Lallensack et al. (2020) whose outline has been represented by 20 landmarks. The three arrows provide examples how the coordinates (*x*, *y*) of landmarks (here shown for the examples 4, 9 and 18) are converted to distances r from the centre. The landmarks shown in red were selected by variable selection via forward step logistic regression. [planned for column width]

**FIGURE 2.**
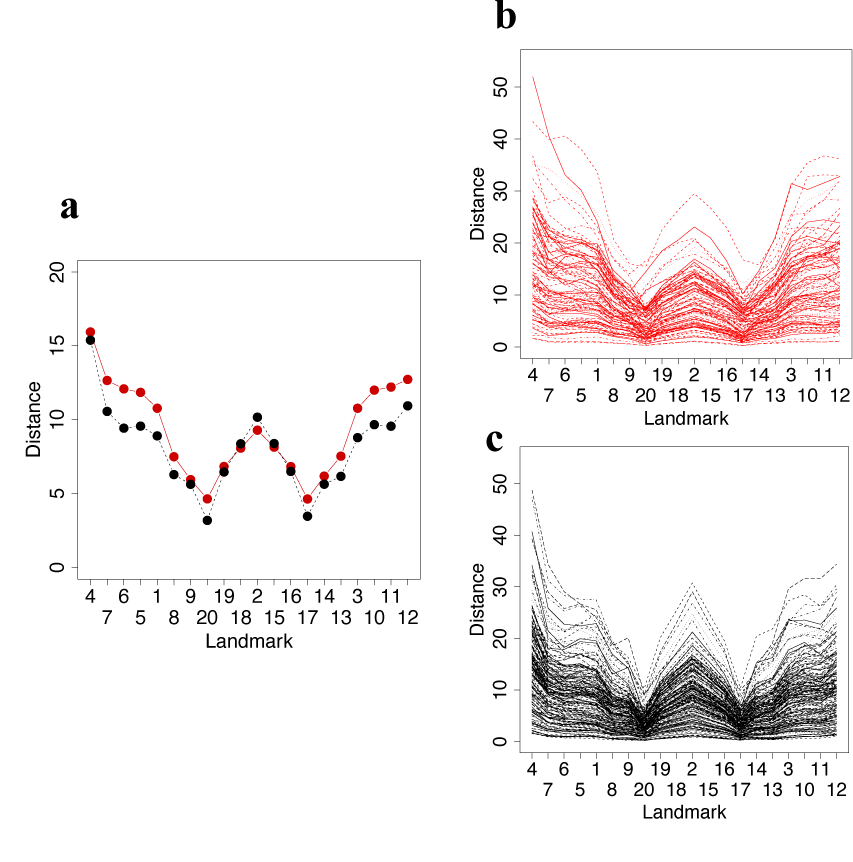
Visualisation of the data set by Lallensack et al. (2020) after transforming the two- dimensional coordinates to distances r from the origin (0,0) according to eq. (1). In the plots, also the sequence of the landmarks is changed, starting with landmark 4 on the left of the x axis and then moving counter-clockwise along the boundary of the footprint. In a, the means of the distances r is plotted for all 20 landmarks for ornithopods (red) and theropods (black). The distances r for all ornithopods are shown in b, theropods are shown in c. [planned for column width]

A slightly more subtle feature of FIGURE 2a relates to landmarks 2, 17 and 20. Here, the distances of landmarks 17 and 20 are again shorter for theropods but on the contrary landmark 2 is on average further away from the centre than the same landmark in ornithopods. Combining these two observations illustrates another distinguishing feature of the two classes, namely that the middle toe of a theropod is usually longer than the middle toe of an ornithopod.

As a whole, all panels in FIGURE 2 illustrate the shape of a three-toed footprint. From landmark 4 which marks the bottom of the footprint, the distance decreases along the right part of the footprint and along the right toe until a minimum is reached at the gap between the right and the middle toe (landmark 20). The distance then increases along the middle toe until the tip is reached (landmark 2). Left of landmark 2 the distance again decreases until the gap between the middle toe and the right toe is reached (landmark 17). Finally, the distance increases again along the left toe and the left side of the footprint. As we can see for both ornothopods (FIGURE 2b) and ornithopods (FIGURE 2c) the graphs for all footprints in the data set follow this pattern although they vary over a wide range – this is related to the footprint size, the distances increase with the footprint size.

The algorithms included in our study were Logistic Regression (LR), Multi-Layer Perceptron (MLP), Random Forest (RF), Support Vector Machine (SVM), Multivariate Adaptive Regression Splines (MARS) and Linear Discriminant Analysis (LDA). More details on these well-known machine learning methods can be found in standard textbooks such as, for example, Hastie et al. (2009). The following software implementations available as packages contained in the programming language `R` (R_Core_Team, 2023) were used: `stats` (LR), `MASS` (LDA), `nnet` (MLP), `randomForestSRC` (RF), `e1071` (SVM), `mda` (MARS).

## RESULTS

The data set, see FIGURE 2, was split into a training set that contained 211 samples (70% of the data) and a test set containing 90 samples (30% of the data) to be used for assessing the performance of the machine learning models used in this study.

For each model, the hyperparameters were optimised to obtain the best possible results. For Logistic Regression (LR), a forward stepwise method was implemented – using the function `step` implemented in R (Hastie & Pregibon, 1992). In Forward Stepwise Regression, landmarks are iteratively added as the performance is assessed via the Akaike Information Criterion (AIC) (Akaike, 1974). The subset of 8 of the 20 landmarks selected in this way were 2, 4, 5, 9, 10, 15, 17, 20, see FIGURE 1.

For the multi-layer perceptron (MLP), the `caret` package was used to carry out a grid search of an MLP implemented in the `nnet` package – the number of neurons in the hidden layer were varied between 1 and 19 and the weight decay parameter λ was changed between 0 and 0.1. Both were optimised using Cohen’s λ as the performance metric (Cohen, 1960). The resulting model had 17 hidden neurons and a weight decay parameter of λ = 0.1. For the Random Forest model, the number of randomly selected predictors was optimised using the `caret` package – the value reached was 4. The optimal SVM with a radial basis function kernel had parameters σ = 0.010, *C* = 32.

After a model is fitted to the training set, its ability to correctly classify theropod and ornithopod footprints is assessed by the relative frequencies of correctly identified theropods (sensitivity) and ornithopods (specificity), respectively. Receiver Operating Characteristic (ROC) curves were generated for each model, see FIGURE 3. The ROC curve illustrates the trade-off of sensitivity and specificity – increasing the performance of a classification method to identify theropods usually comes at the expense of its ability to recognise ornithopods and vice versa. By changing the threshold that determines which samples are classified as theropods versus ornithopods, sensitivity and specificity of a classification method can be varied. In FIGURE 3 we have indicated by red points sensitivity and specificity that can be achieved for a given method by optimising the threshold. A summary statistic for ROC curves is the area under the ROC curve (AUROC) which provides a metric of the overall performance of a classifier.

**FIGURE 3.**
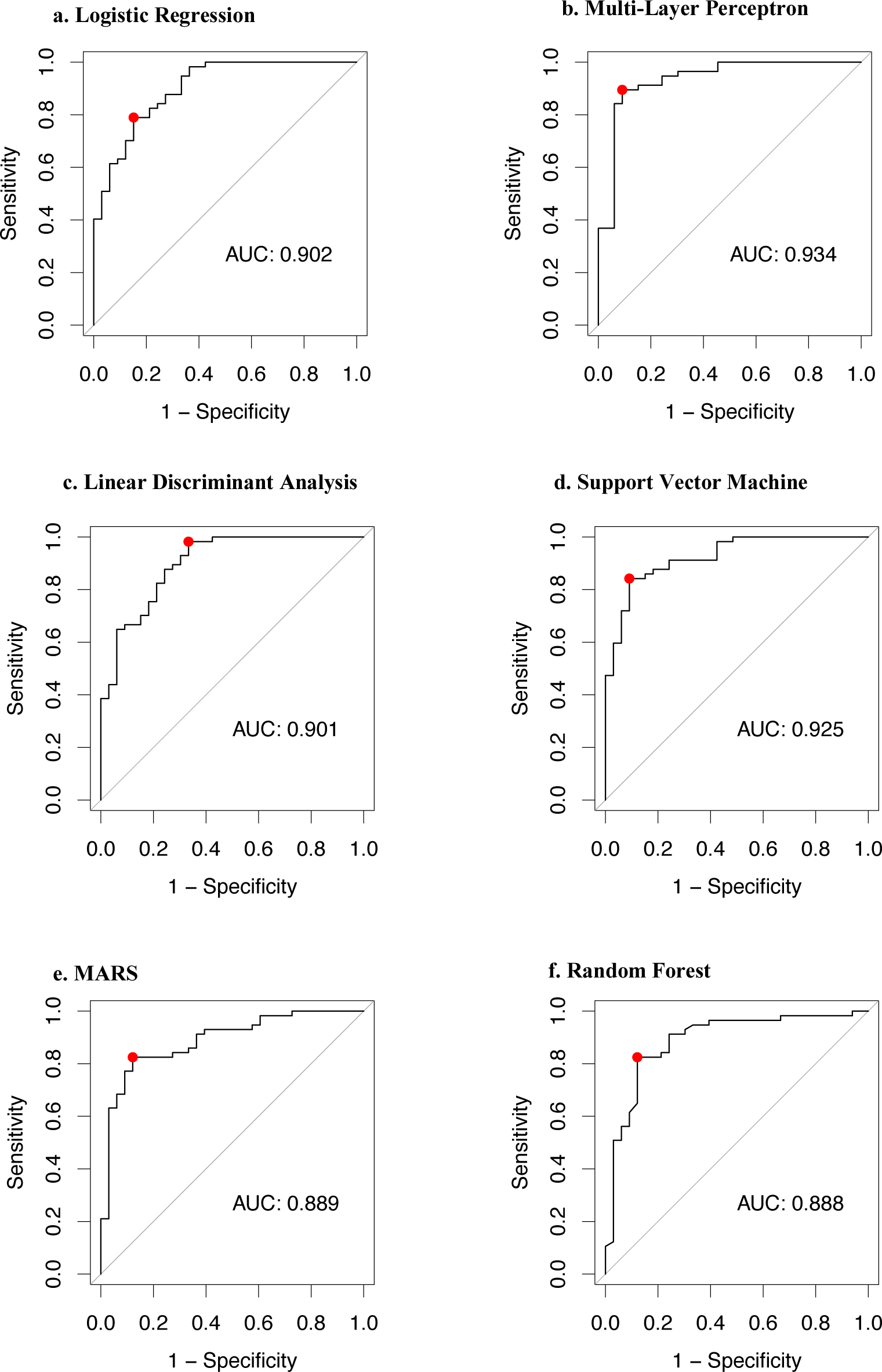
ROC curves for the machine learning methods used in this study. For each method, the Area under the Curve (AUC) has been calculated. The red points show the recall of theropods (sensitivity) and ornithopods (specificity) achieved by the optimal threshold. [planned for page width]

TABLE 1 shows confusion matrices and performance metrics, FIGURE 4 shows the recall and accuracy for all methods both before and after optimising the threshold using the ROC curves. All models show good accuracy scores between 80% and 90%. However, all methods except MLP are notably worse at accurately predicting the smaller class i.e. ornithopods before the threshold of the classifier is adjusted. After moving the threshold, most methods increase the recall of ornithopods whilst decreasing the recall of theropods. This is expected because there is a trade-off between both metrics which is represented by the ROC curve. An exception to this pattern is LDA. Before adjusting the threshold, LDA has the best recall for theropods of any of the 6 methods (91.2%) whilst at the same time having the worst recall for ornithopods (69.7%). After adjusting the threshold, the recall of theropods increases whereas the recall of ornithopods decreases even further – as a result LDA becomes an extreme case of a classifier that reaches a recall of 98.2% for theropods but at the expense of a very weak performance of classifying ornithopods, achieving a recall of only 66.7%. In contrast, even before adjusting the threshold, MLP’s recall of ornithopods is already 90.9%, which remains unchanged after moving the threshold whilst the recall of theropods increases from 87.7% to 89.5%. Thus, MLP is not only the method that achieves the highest accuracy (90%) but also the highest recall for both classes. Because for this study performance should ideally be as high as possible for classifying both ornithopods and theropods, the MLP is superior to the other models.

**TABLE 1.**
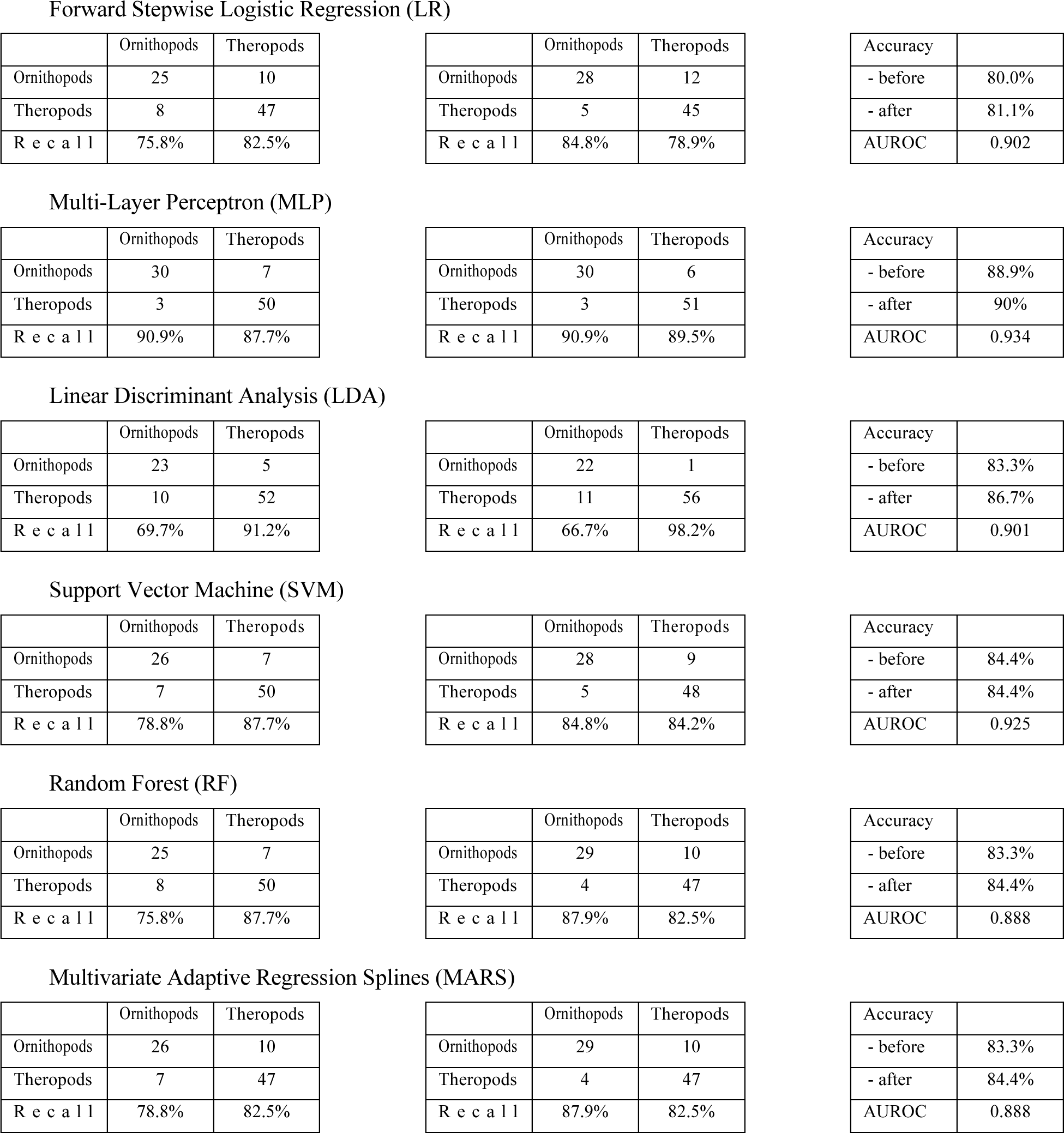
Confusion matrices with performance metrics for several machine learning methods. Recall of ornithopods and theropods quantifies the percentages of correctly classified samples of each type of dinosaur whereas accuracy is the overall percentage of correctly classified footprints. All these metrics were calculated before and after adjusting the thresholds from 0.5 as indicated by the red points in Figure 3. Finally, AUROC is reported for all methods. Plots of recall and accuracy for all methods before and after adjusting the thresholds are shown in Figure 4.

**FIGURE 4.**
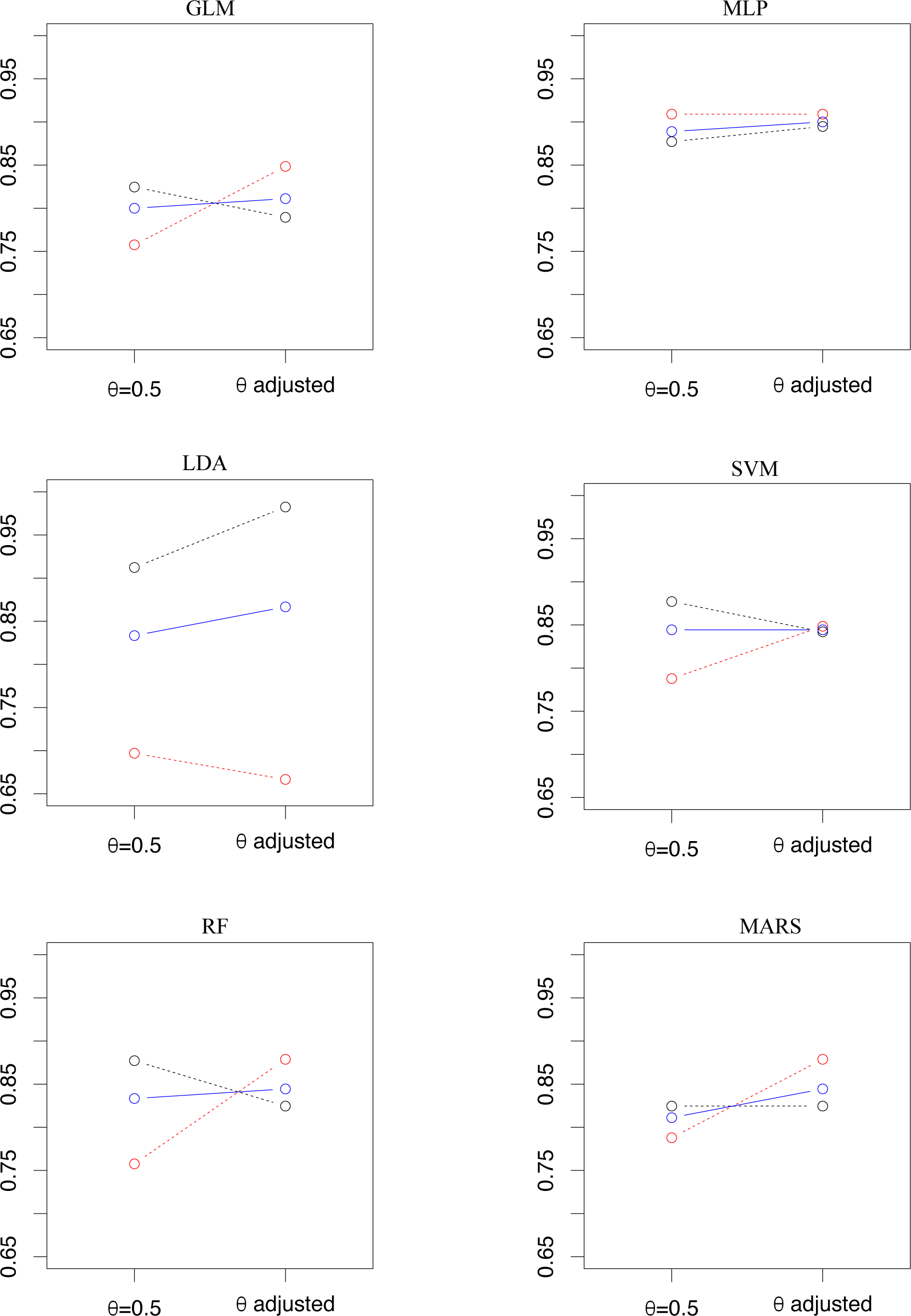
The plots show recall of ornithopods (red, dashed) and theropods (black, dashed) as well as accuracy (blue, solid) for all methods before and after adjusting the threshold of the classifier. Recall quantifies the proportion of correctly classified samples of each type of dinosaur whereas accuracy is the overall percentage of correctly classified footprints. [planned for page width]

## DISCUSSION

Six machine learning methods were applied to the problem of distinguishing theropods and ornithopods, two groups of dinosaurs, based on footprints. Each footprint is represented by 20 landmarks which are placed as explained by Lallensack et al. (2020). Of the six models, MLP emerges as clearly superior in comparison to the other methods, achieving a recall for both classes of about 90%.

One feature of the approach presented here is that it relies on the system of 20 landmarks proposed by Lallensack et al. (2020), a set of points of interest located on the boundaries of the footprints, such as the tips of the three toes (landmarks 1, 2 and 3) or the rear (landmark 4) of each footprint. One immediate advantage of using landmarks rather than working directly with image data is robustness to sources of error such as, for example, footprints that are located off-centre or are rotated whilst, when determining the landmarks from images, distortions such as different orientations of a footprint with respect to the image frame are removed. As a result, using landmarks for building classifiers for distinguishing between theropods and ornithopods does not need to address the usual challenges faced in image processing applications. However, this suggests that the accuracy that can be achieved using machine learning models depends on the quality of the landmark data set.

Instead of using the two-dimensional coordinates (*x*, *y*) that indicate the locations of each landmark, we instead work with the distances 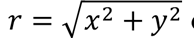 of the landmarks from the centre of the footprint. Thus, rather than using 2 × 20 = 40 coordinates, we reduce the dimensionality of the data to 20 distances from the centre. From FIGURE 2 it seems intuitively clear that not much information is lost due to this transformation of the data set. This can be explained more rigorously by considering that rather than by two-dimensional cartesian coordinates (*x*, *y*), the locations of landmarks can equivalently be represented by polar coordinates (*r*, *ϕ*) i.e. by the distance *r* from the origin and an angle *ϕ* that indicates the direction where a point is located. Thus, our transformed data set can be described as a transformation to polar coordinates where the angular variable *ϕ* is omitted. Each landmark lies in a particular direction from the centre – landmark 2, the tip of the middle toe, lies above the centre, whereas landmark 4 is located below. Landmarks 1 is lateral (on the upper right), and landmark 2 is medial (on the upper left). Thus, because for different footprints each landmark is expected to be located in a similar direction from the centre most of the information is contained in the distance *r* from the centre and the angle *ϕ* can be omitted.

In fact, replacing the cartesian coordinates (*x*, *y*) by distances *r* from the centre is essential for obtaining a set of variables that is suitable for training machine learning methods. When attempting to train various machine learning methods on the original data set, the fact that some coordinates are not informative – for example, the *x* coordinate of landmark 2 is close to zero whereas the *y* coordinates of landmarks 5, 8, 10, 13, 17 and 20 are small – causes problems and it is difficult to achieve convergence.

The number of variables can be reduced even further by variable selection. This was implemented for Logistic Regression via Forward Step Regression. The landmarks selected by forward regression – shown in red in FIGURE 3 – were among those that would intuitively be expected to be in the subset of the most informative landmarks – landmarks 2 and 4 as the tip and rear of the footprint which are related the length of a footprint, 5 and 10 that are linked to the width of the footprint, 17 and 20 halfway along the middle toe, and 9 at the beginning of the lateral (right) toe. Only landmark 15 seems slightly inconsistent with this pattern because it is the landmark next to landmark 2 – intuitively one might rather expect to find 14, the landmark opposite of 9, in the set of selected variables instead.

When fitting the other machine learning methods to the subset of variables selected by forward regression, the models obtained showed similar performance as when fitted to the full data set. Arguably that suggests that more parsimonious models could be obtained when reducing the modelling to the landmarks selected by forward selection.

Finally, it is interesting to investigate which footprints were misclassified in order to explore possible limitations of the approach, see TABLE 2. Only two ornithopod and one theropod footprint were misclassified by all six machine learning models, see FIGURE 5a. The shapes of these three footprints, published by Castanera et al. (2013), Li et al. (2012) and Xing, Lockley, Zhang, et al. (2014), reveal why it might be challenging to classify them correctly – for example, in all three figures the middle toe clearly points to the left or to the right, respectively, rather than straight ahead. Indeed, Castanera et al. (2013) argued that the tracks they described are challenging to refer to either group, and in this case the assignment to ornithopods could be made only because of the presence of manus tracks, though other works have attributed manus tracks to theropod trackmakers (e.g. Milner et al. (2009); Li et al. (2019)). The Li et al. (2012) track is referred to the ichnogenus *Anomoepus*, which is produced by small, basal ornithischians. In the absence of manus tracks, such tracks are generally difficult to assign to either theropods or ornithischians, and it is possible that the Li et al. (2012) track is a mis-identified theropod track.

**TABLE 2.** Samples in the data set that have been misclassified by one or more of the methods after adjusting the threshold. References to all sources can be found in Lallensack et al. (2020).

| Source | Ichnogenus | Epoch | Group | lda | glm | nnet | rf | svm | mars | nMisclass |
| --- | --- | --- | --- | --- | --- | --- | --- | --- | --- | --- |
| barco 2005 1 | *Iberosauripus* | Cretaceous | Cretaceous Early | ✓ | ✓ | ✓ | ✓ | ✓ | ✗ | 1 |
| castanera 2013b 1 | NA | NA | NA | ✗ | ✗ | ✗ | ✗ | ✗ | ✗ | 6 |
| castanera 2013b 2 | NA | NA | NA | ✗ | ✗ | ✓ | ✗ | ✗ | ✗ | 5 |
| dalman 2013 1 | *Anomoepus* | Jurassic | Jurassic Early | ✗ | ✓ | ✗ | ✗ | ✓ | ✗ | 4 |
| ellenberger 1974 2 | *Moyenisauropus* | Jurassic | Jurassic Early | ✓ | ✓ | ✓ | ✓ | ✓ | ✗ | 1 |
| ellenberger 1974 3 | *Moyenisauropus* | Jurassic | Jurassic Early | ✗ | ✗ | ✓ | ✓ | ✗ | ✓ | 3 |
| ellenberger 1974 5 | *Moyenisauropus* | Jurassic | Jurassic Early | ✗ | ✗ | ✓ | ✓ | ✗ | ✓ | 3 |
| gierlinski 2004 6 | *Anomoepus* | Jurassic | Jurassic Early | ✗ | ✗ | ✓ | ✓ | ✗ | ✓ | 3 |
| gierlinski 2009b 3 | *Therangospodus* | Jurassic | Jurassic Middle | ✓ | ✗ | ✓ | ✓ | ✗ | ✓ | 2 |
| gierlinski 2009b 4 | *Carmelopodus* | Jurassic | Jurassic Middle | ✓ | ✓ | ✗ | ✗ | ✗ | ✗ | 4 |
| jianjun 2012 3 | *Anomoepus* | Jurassic | NA | ✗ | ✗ | ✗ | ✗ | ✗ | ✗ | 6 |
| kim 2017 3 | *Corpulentatpus* | Cretaceous | Cretaceous Early | ✓ | ✗ | ✗ | ✗ | ✗ | ✗ | 5 |
| lallensack 2016 2 | NA | Cretaceous | Cretaceous Early | ✓ | ✗ | ✗ | ✓ | ✗ | ✗ | 4 |
| lockley 1998b 2 | *Megalosauripus* | Jurassic | Jurassic Late | ✓ | ✓ | ✓ | ✗ | ✓ | ✗ | 2 |
| lockley 1998c 5 | *Therangospodus* | Jurassic | Jurassic Late | ✓ | ✓ | ✗ | ✓ | ✓ | ✓ | 1 |
| lockley 2008 4 | *Hispanosauropus* | Jurassic | Jurassic Late | ✓ | ✓ | ✓ | ✗ | ✓ | ✓ | 1 |
| lockley 2008b 1 | *Minisauripus* | Cretaceous | Cretaceous Late | ✗ | ✗ | ✓ | ✓ | ✗ | ✓ | 3 |
| lockley 2008b 3 | *Minisauripus* | Cretaceous | Cretaceous Early | ✓ | ✗ | ✓ | ✓ | ✗ | ✓ | 2 |
| lockley 2009 4 | *Dinehichnus* | Jurassic | Jurassic Early | ✗ | ✗ | ✓ | ✗ | ✗ | ✗ | 5 |
| lockley 2009 5 | *Dinehichnus* | Jurassic | Jurassic Early | ✗ | ✓ | ✓ | ✓ | ✗ | ✓ | 2 |
| lockley 2014b 1 | *Irenesauripus* | Cretaceous | NA | ✓ | ✓ | ✓ | ✗ | ✓ | ✓ | 1 |
| lockley 2014e 1 | NA | Cretaceous | NA | ✓ | ✓ | ✓ | ✗ | ✓ | ✓ | 1 |
| matsukawa 2006 6 | NA | Cretaceous | Cretaceous Early | ✗ | ✓ | ✓ | ✓ | ✓ | ✓ | 1 |
| olsen 1980 2 | *Grallator* | Jurassic | Jurassic Early | ✓ | ✗ | ✓ | ✓ | ✗ | ✓ | 2 |
| olsen 2003 3 | *Anomoepus* | Jurassic | Jurassic Early | ✗ | ✗ | ✓ | ✗ | ✗ | ✗ | 5 |
| pittman 1989 3 | NA | Cretaceous | Cretaceous Early | ✓ | ✓ | ✓ | ✓ | ✓ | ✗ | 1 |
| raath 1972 1 | NA | NA | NA | ✗ | ✗ | ✓ | ✓ | ✗ | ✗ | 4 |
| xing 2011 4 | *Kayentapus* | Cretaceous | Cretaceous Early | ✗ | ✗ | ✗ | ✓ | ✗ | ✗ | 5 |
| xing 2014f 1 | *Paracorpulentapus* | Cretaceous | Cretaceous Late | ✗ | ✗ | ✗ | ✗ | ✗ | ✗ | 6 |
| xing 2014h 1 | NA | Jurassic | Jurassic Early | ✓ | ✓ | ✗ | ✗ | ✗ | ✗ | 4 |
| xing 2016c 1 | *Anomoepus* | Jurassic | Jurassic Early | ✓ | ✓ | ✓ | ✗ | ✓ | ✓ | 1 |
| xing 2016g 1 | *Minisauripus* | Cretaceous | Cretaceous Early | ✓ | ✗ | ✓ | ✓ | ✗ | ✓ | 2 |

**FIGURE 5.**
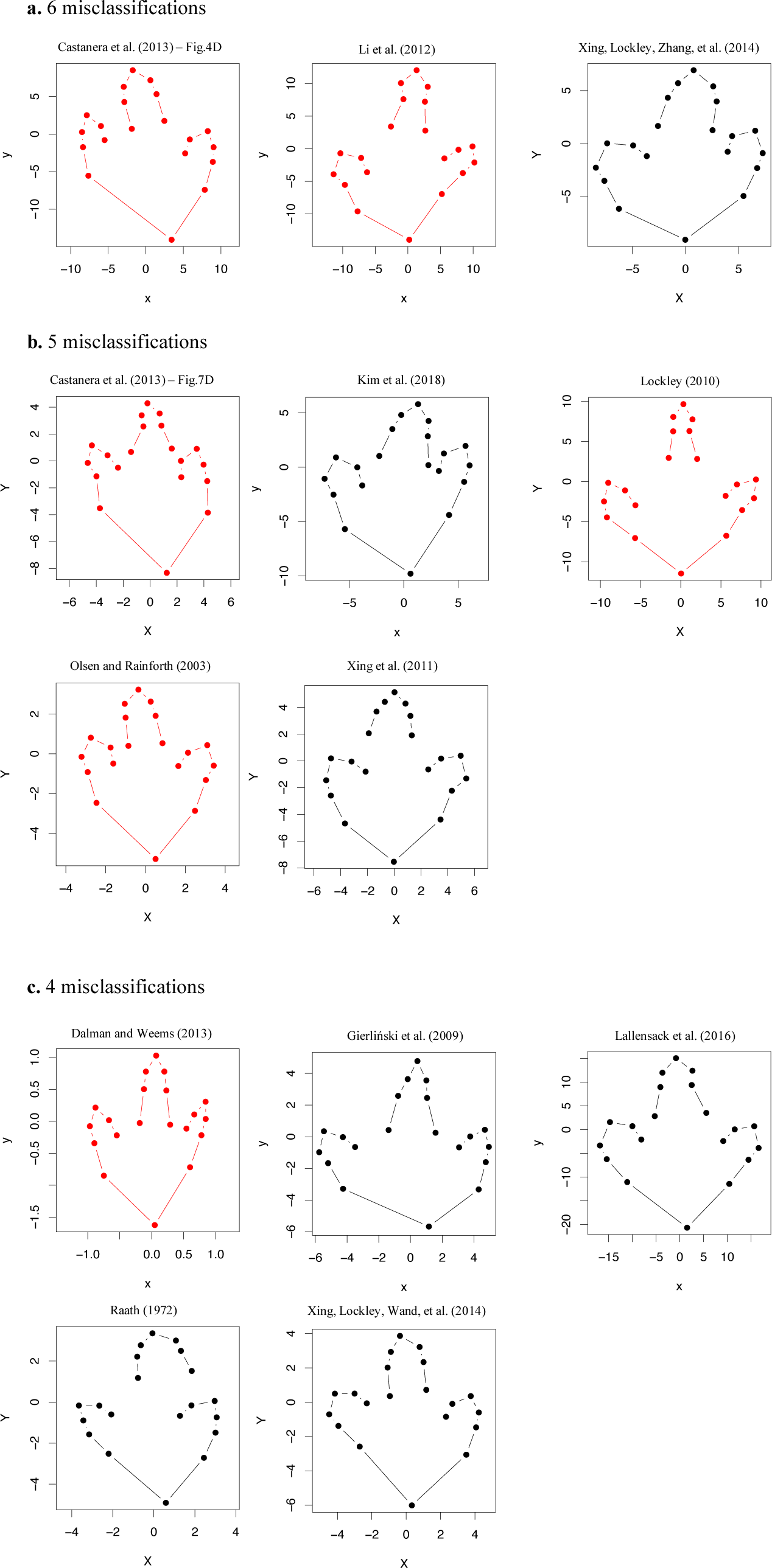
Footprints that are misclassified by all (6), 5 or 4 of the methods applied in this study. Ornithopod footprints are shown in red, theropods in black. Note that the footprints shown here have different sizes – they are plotted on different scales to emphasise their shapes rather than their sizes. [planned for page width]

The five footprints that have been misclassified by 5 out of 6 methods have been published in Castanera et al. (2013), Kim et al. (2018), Lockley (2010); Olsen and Rainforth (2003) and Xing et al. (2011). Looking at other misclassified footprints, for example, those shown in FIGURE 5c, it appears that the theropod footprints that have been misclassified by 4 out of 6 machine learning methods show characteristics that are more common for ornithopods – for Dalman and Weems (2013), Gierliński et al. (2009), Lallensack et al. (2016), Raath (1972) Xing, Lockley, Wand, et al. (2014) the middle toe is relatively short and wide rather than long and thin. Again, some theropod tracks are very ornithopod-like in appearance (see Milner et al. (2023) for a particularly deceiving example). Consequently, some ambiguity is expected, and it is likely that some of the tracks are impossible to be classified correctly.

## CONCLUSION

We developed a machine learning approach for classifying ornithopod and theropod footprints using a data set consisting of 20 landmarks that describe the shape of the footprints. We achieve accuracies of above 80% up to 90% for MLP, the best-performing method. MLP also shows a balanced performance for both classes, achieving recalls of approximately 90% for both ornithopods and theropods.

## Supporting information

Metadata for footprints

Data set

Knitted R Markdown report

## ACKNOWLEDGMENTS

JNL was supported by the Alexander von Humboldt-Foundation, Bonn, Germany. PLF was supported by a UKRI Frontier Research Grant TRACKEVOL (selected by the ERC as a consolidator award).

## AUTHOR CONTRIBUTIONS

IS, JNL, PLF and IJ designed the project. MJ and IS carried out the project and wrote the first draft of the manuscript. All authors reviewed and approved the manuscript.

## SUPPLEMENTARY FILE(S)

- JonesEtAl.Rmd: `R Markdown` implementation of the machine learning methods used for this study.
- JonesEtAl.PDF: Report generated from `R Markdown` file.
- DinoPrints.csv: Data set of ornithopod and theropod footprints (Lallensack et al., 2020).

## Notes

https://github.com/merlinthemagician/dinoprints-ML

### Competing Interest Statement

The authors have declared no competing interest.

