## Supplementary material for "Classification of dinosaur footprints using machine learning": Knitted R Markdown report

### Dinosaur footprints landmarks data set

The data set consists of landmarks that represent dinosaur footprints. The variables `X` and `Y` are the coordinates of the landmarks, `id` assigns a number to each footprint and `Group` is the group of dinosaurs that a footprint belongs to:

```
dinoprints <- read.csv("DinoPrints.csv", stringsAsFactors = TRUE)
knitr::kable(head(dinoprints))
```

| X | Y | id | Group |
|---:|---:|---:|---|
| 22.2674712 | 5.265072 | 1 | Theropoda |
| 0.0956471 | 21.257381 | 1 | Theropoda |
| -22.6655232 | 5.895802 | 1 | Theropoda |
| 6.0806097 | -40.237751 | 1 | Theropoda |
| 22.3706016 | -1.401412 | 1 | Theropoda |
| 21.3634667 | -7.817616 | 1 | Theropoda |

As an example, we plot the first three footprints.

```
par(pty="s")
plot(subset(dinoprints, id==1, select = c(X, Y)),
     pch = 19, col="darkgreen",
     asp=1)

## I have added two more footprints to the plot
points(subset(dinoprints, id==2, select = c(X, Y)),
     pch = 19, col="red")

points(subset(dinoprints, id==3, select = c(X, Y)),
     pch = 19, col="blue")
```

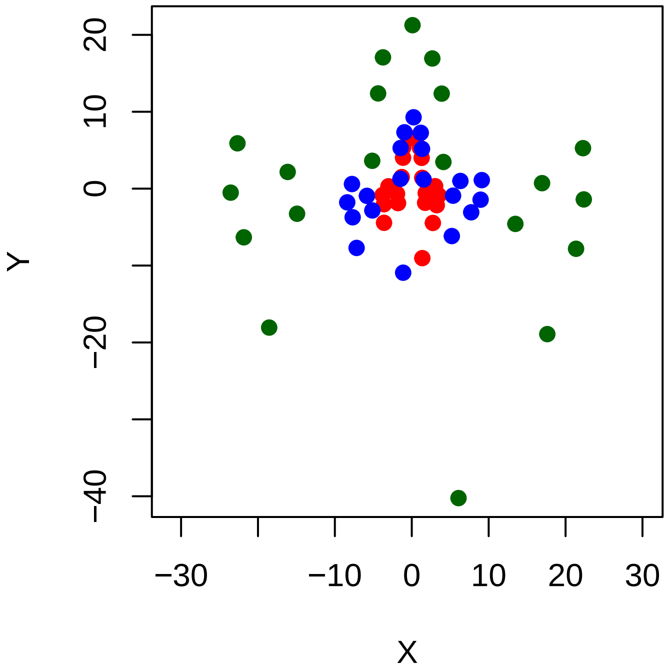


### Transforming the data set

For each footprint (i.e. all 20 `X` and `Y` coordinates for a given 'id) we transform the coordinates to lengths $r$:

$$r = \sqrt{x^2 + y^2}$$

These are the distances from the centre $(0, 0)$ of the footprint.

We genereate a new data frame for which each row contains the 20 distances $r_1, \ldots, r_{20}$ and the group `Group` that the dinosaur belongs to.

#### One footprint

This is how a single footprint can be transformed - we select the first footprint and calculate the distance $r$ for each landmark:

```
footNo1 <- subset(dinoprints, id==1, select=c(X,Y))
sqrt(footNo1$X^2 + footNo1$Y^2)
```

```
##  [1] 22.881461 21.257596 23.419787 40.694599 22.414455 22.748908 25.855709
##  [8] 16.960379 14.231144 23.554969 22.736981 25.887193 16.279754 15.268953
## [15] 17.477585 13.133103  6.289382 17.138873 12.956998  5.382066
```

#### All footprints

We now calculate distances from the centre for all footprints:

```
dino.dist <- data.frame(matrix(nrow = max(dinoprints$id), ncol = 20))
nDinos <- nrow(dino.dist)
for(k in 1:nDinos) {
  footprint <- subset(dinoprints, id==k, select=c(X,Y))
  dino.dist[k, ] <- sqrt(footprint$X^2 + footprint$Y^2)
}
```

```r
#knitr::kable(head(dino.dist)) %>% kable_styling(latex_options = "scale_down")
pander(head(dino.dist), split.table = 80, style = 'rmarkdown', position="left")
```

Table 1: Table continues below

| X1 | X2 | X3 | X4 | X5 | X6 | X7 | X8 | X9 |
|---|---|---|---|---|---|---|---|---|
| 22.88 | 21.26 | 23.42 | 40.69 | 22.41 | 22.75 | 25.86 | 16.96 | 14.23 |
| 3.055 | 6.571 | 3.045 | 9.137 | 3.611 | 3.897 | 5.233 | 1.905 | 2.532 |
| 9.174 | 9.284 | 7.798 | 10.99 | 9.065 | 8.318 | 8.071 | 6.403 | 5.446 |
| 23.15 | 23.6 | 23.18 | 39.34 | 26.39 | 25.31 | 27.9 | 16.21 | 14.13 |
| 10.24 | 11.57 | 9.878 | 18.47 | 10.91 | 11.08 | 12.73 | 7.145 | 7.364 |
| 9.073 | 8.954 | 7.156 | 14.47 | 9.202 | 9.189 | 9.611 | 6.807 | 5.614 |

Table 2: Table continues below

| X10 | X11 | X12 | X13 | X14 | X15 | X16 | X17 | X18 |
|---|---|---|---|---|---|---|---|---|
| 23.55 | 22.74 | 25.89 | 16.28 | 15.27 | 17.48 | 13.13 | 6.289 | 17.14 |
| 3.888 | 4.141 | 5.721 | 2.029 | 2.591 | 5.473 | 4.18 | 1.997 | 5.401 |
| 8.578 | 8.542 | 10.53 | 5.904 | 5.858 | 7.391 | 5.485 | 1.947 | 7.352 |
| 26.53 | 26.35 | 30.39 | 15.62 | 14.44 | 18.32 | 13.77 | 8.53 | 18.75 |
| 10.79 | 10.64 | 12.23 | 7.049 | 6.704 | 9.367 | 7.514 | 4.607 | 10 |
| 8.355 | 8.268 | 9.825 | 5.023 | 4.585 | 7.379 | 5.878 | 3.184 | 7.136 |

| X19 | X20 |
|---|---|
| 12.96 | 5.382 |
| 4.204 | 1.957 |
| 5.339 | 1.946 |
| 13.12 | 9.12 |
| 8.045 | 3.858 |
| 5.478 | 2.403 |

#### Finding the `Group` label for each sample

Now, the `Group` label for each sample is added.

```r
# If we start dino.group as a vector, the factor structure will be destroyed.
dino.group<-factor()
for(k in 1:nrow(dino.dist)) {
  footGroup <- subset(dinoprints, id==k, select=Group)
  # We take the first entry of Group for each footprint (they should be all the same)
  # and append them to dino.group
  dino.group <- c(dino.group, footGroup$Group[1])
}

dino.dist$Group <- dino.group
```

### Visualising distances from the origin

First, we plot the first footprint showing the indices of each landmark next to the points.

```
# Make square-shaped plot
par( pty="s")

plot(subset(dinoprints, id==1, select = c(X, Y)),
     xlim = c(-25,25), ylim=c(-45,25),
     pch=19, col="darkgreen",
     #main = "Footprint 1",
     xlab="x", ylab = "y", asp=1)

## Add labels next to points
for (i in 1:20) {
  text(dinoprints[i,"X"]+2, dinoprints[i,"Y"]+2, labels = i,
       cex=0.75, col="black")
}
```

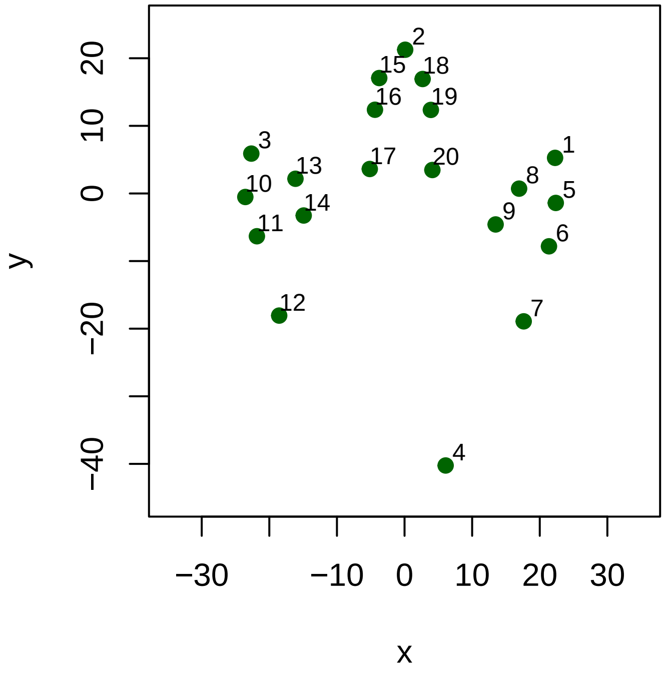


In order to visualise the transformed data we rearrange the landmarks starting from landmark 4 which marks the base of the footprint and then rotating around the boundary of the footprint in anti-clockwise direction. This enables us to show the variation of distances from the centre as we move around the boundary of the footprint. We show the mean distances from the centre:

```
lmCircle <- c(4,7,6,5,1,8,9,20,19,18,2,15,16,17,14,13,3,10,11,12)

landmarkLab <- "Landmark"
distLab <- "Distance"

dino.dist.split <- split(subset(dino.dist, select=-Group), dino.dist$Group)
dino.dist.means <- lapply(dino.dist.split, colMeans)

matplot(
  cbind(dino.dist.means$Ornithischia[lmCircle],
        dino.dist.means$Theropoda[lmCircle]
  ), col=c("red3", "black"), type="b", pch=19, ylim=c(0,20)
  ,xlab=landmarkLab, ylab=distLab,
```

```
  xaxt="n",
  main = "Mean distances"
)
axis(1,at=1:length(lmCircle),labels = FALSE, cex.axis=0.5)
for(i in 1:length(lmCircle)) mtext(1, at=i, text=lmCircle[i],line=((i+1)%%2)*1+0.5)
```

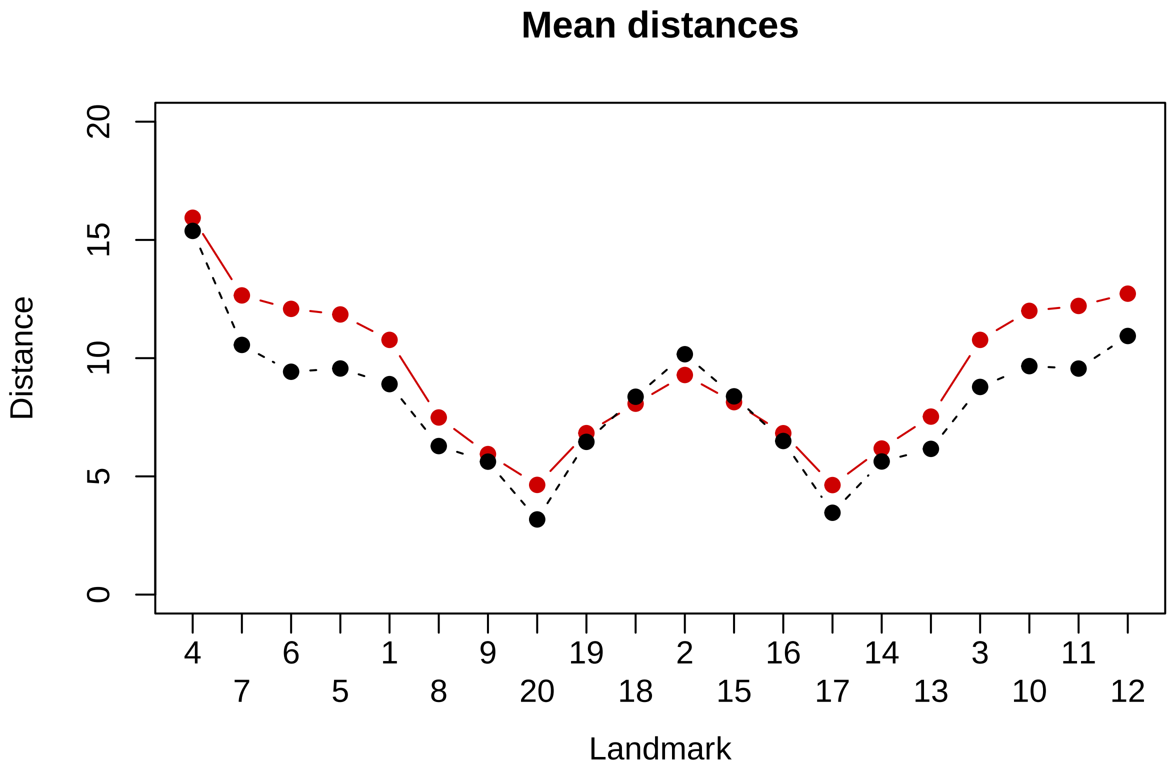


We also plot the median distances:

```
library(robustbase)

dino.dist.medians <- lapply(dino.dist.split, function(x) colMedians(as.matrix(x)))

matplot(
  cbind(dino.dist.medians$Ornithischia[lmCircle],
        dino.dist.medians$Theropoda[lmCircle]
  ), col=c("red3", "black"), type="b", pch=19, ylim=c(0,20)
  ,xlab=landmarkLab, ylab=distLab,
  xaxt="n",
  main = "Median distances"
)
axis(1,at=1:length(lmCircle),labels = FALSE, cex.axis=0.5)
for(i in 1:length(lmCircle)) mtext(1, at=i, text=lmCircle[i],line=((i+1)%%2)*1+0.5)
```

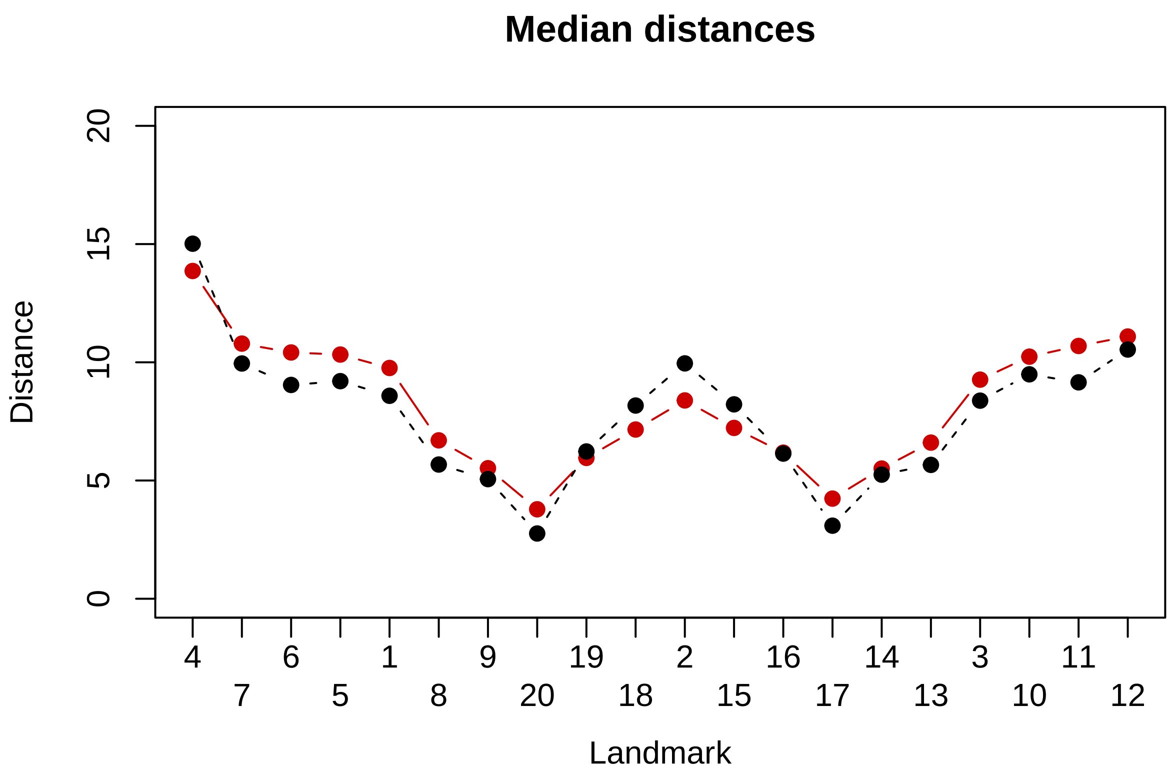


We plot the theropods in black:

```
matplot(t(subset(dino.dist,
                 subset=Group=="Theropoda"))[lmCircle,], type="l", col="black",
        ylim=c(0,55), ylab=distLab,
        xaxt="n")
axis(1,at=1:length(lmCircle),labels = FALSE, cex.axis=1)
for(i in 1:length(lmCircle)) {
  mtext(1, at=i, text=lmCircle[i],line=((i+1)%%2)*1.2+1, cex = 1)
}
mtext(1, adj=0.5, text=landmarkLab, line = 3.5, cex=1)
```

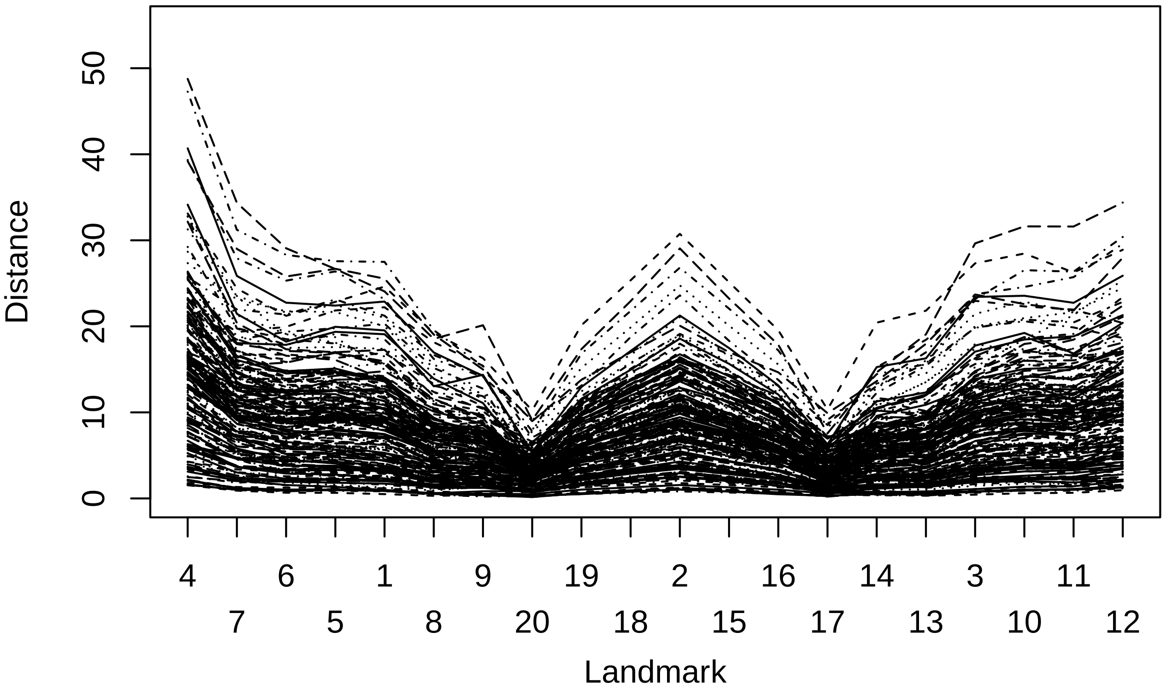


This plot shows the ornithopods in red:

```
matplot(t(subset(dino.dist,
                 subset=Group=="Ornithischia"))[lmCircle,], type="l", col="red",
        ylim=c(0,55), ylab=distLab,
        xaxt="n")
axis(1,at=1:length(lmCircle),labels = FALSE, cex.axis=1)
for(i in 1:length(lmCircle)) {
  mtext(1, at=i, text=lmCircle[i],line=((i+1)%%2)*1.2+1, cex = 1)
}
mtext(1, adj=0.5, text=landmarkLab, line = 3.5, cex=1)
```

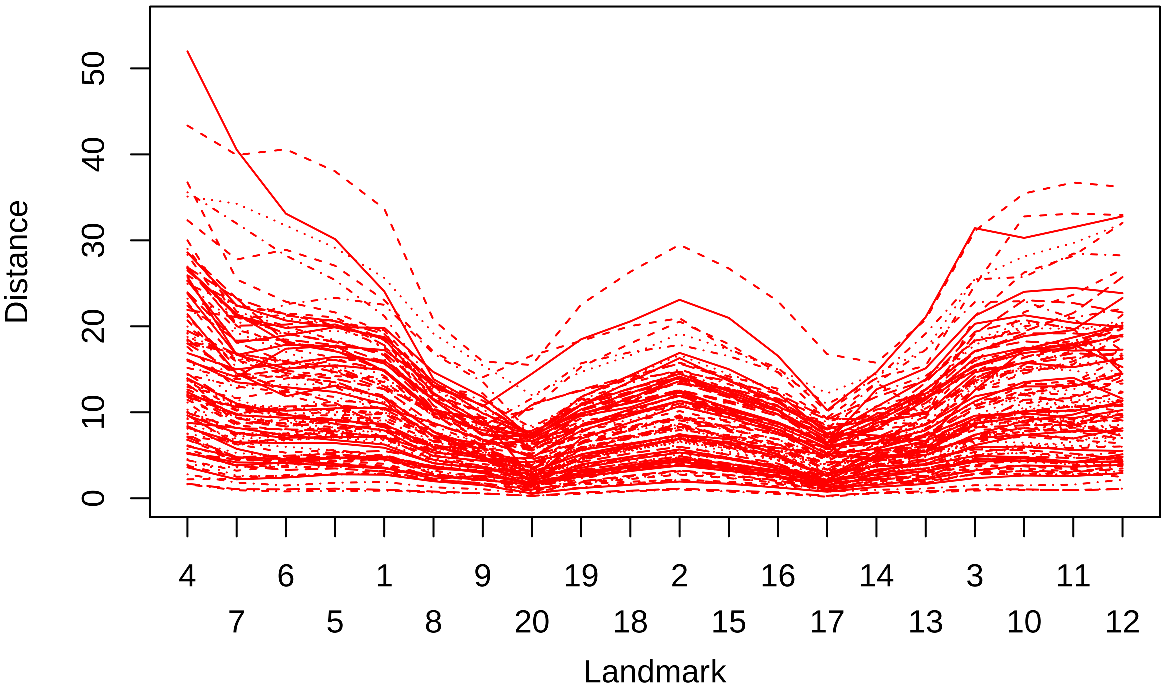


### Machine learning

After representing each footprint via the distances of 20 landmarks to the centre of the footprints we train five methods - Linear Discriminant Analysis (LDA), Logistic Regression (LR), Multi-layer perceptron (MLP), Random Forest (RF), Support Vector Machine (SVM) and Multivariate Adaptive Regression Splines (MARS) - for distinguishing *Theropoda* and *Ornithischia*.

#### Training/testing split

All methods will be trained on data using the same train/test split.

```
set.seed(10)

trainTestRatio <- 0.7
spt <- sample(c(rep(0, trainTestRatio * nrow(dino.dist)),
                rep(1, (1 - trainTestRatio) * nrow(dino.dist))))
train.dino.dist <- dino.dist[spt == 0,]
test.dino.dist <- dino.dist[spt == 1,]
```

Train/test ratio is set to 70%.

```
# Calculate sensitivity, specificity and accuracy for a confusion matrix.
confMatStats <- function(confMat) {
  specificity <- confMat[1,1]/sum(confMat[,1])
  sensitivity <- confMat[2,2]/sum(confMat[,2])
  accuracy <- sum(diag(confMat))/sum(confMat)

  return(
    c(Sensitivity=sensitivity, Specificity=specificity, Accuracy=accuracy)
  )

}
```

All classification methods are trained using the training data set. The classification threshold is adjusted using the receiving operator characteristic (ROC) curve which compensates for the influence of the slight imbalance - the data contain more *Theropoda* than *Ornithischia* samples.

#### Linear Discriminant Analysis (LDA)

Results before adjusting the threshold.

```
library(MASS)
library(pROC)

lda <- lda(Group ~. , data=train.dino.dist)
lda.predict<- predict(lda, newdata = test.dino.dist)
ldaConf<-table(
  ifelse(lda.predict$posterior[,2]<0.5,
         levels(test.dino.dist$Group)[1],
         levels(test.dino.dist$Group)[2]), test.dino.dist$Group
  )
knitr::kable(ldaConf)
```

| | Ornithischia | Theropoda |
|---|---|---|
| Ornithischia | 23 | 5 |
| Theropoda | 10 | 52 |

```
stats.lda <- confMatStats(ldaConf)
knitr::kable(stats.lda)
```

| | x |
|---|---|
| Sensitivity | 0.9122807 |
| Specificity | 0.6969697 |
| Accuracy | 0.8333333 |

Determining an adjusted threshold via the ROC curve.

```
par(pty="s")
lda.roc <- roc(test.dino.dist$Group, lda.predict$posterior[,2],
               print.auc=TRUE,
               # auc.polygon=TRUE,
               plot=TRUE,
               print.auc.x=0.5,print.auc.y=0.3,
               legacy.axes=TRUE,
               xlim=c(1,0), ylim=c(0,1),
               asp=1
               #,main="Linear Discriminant Analysis (LDA)"
               )
lda.Opt <- coords(lda.roc, "best")

points(lda.Opt$specificity, lda.Opt$sensitivity, col="red", pch=19, cex=1.5)
```

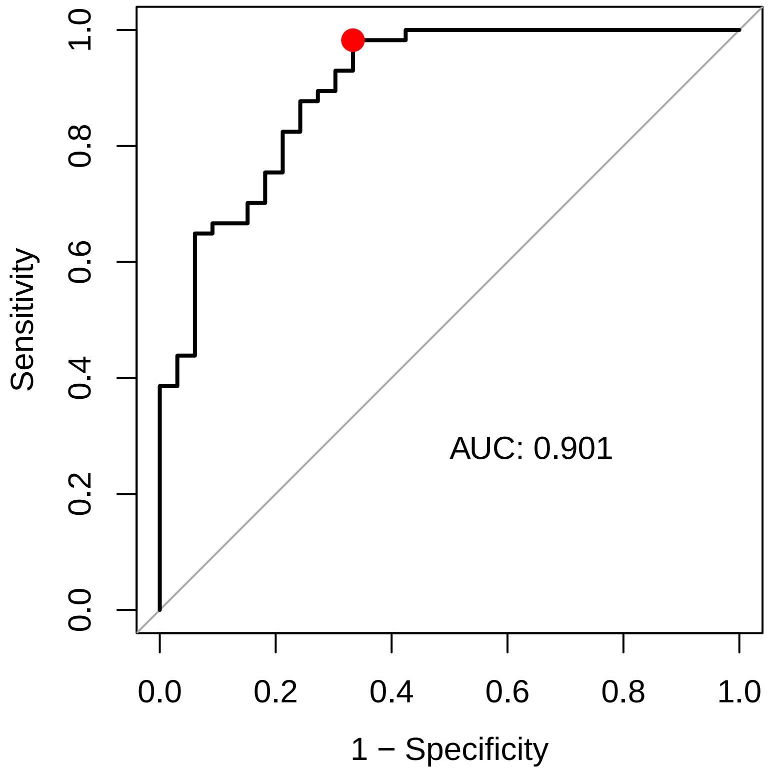


```
if(plotPDF) {
  pdf("figures/LDA_ROC.pdf")
  par(mar=c(4,4,1,1), cex=2)
  lda.roc <- roc(test.dino.dist$Group, lda.predict$posterior[,2],
                print.auc=TRUE,
                # auc.polygon=TRUE,
                plot=TRUE,
                print.auc.x=0.5,print.auc.y=0.3,
                print.auc.cex=1,
                legacy.axes=TRUE,
                xlim=c(1,0), ylim=c(0,1),
                asp=1
               #,main="Linear Discriminant Analysis (LDA)"
              )
  lda.Opt <- coords(lda.roc, "best")

  points(lda.Opt$specificity, lda.Opt$sensitivity, col="red", pch=19, cex=1)
  dev.off()
}
```

Results after adjusting the threshold.

```
ldaConf.Opt<-table(
  ifelse(lda.predict$posterior[,2]<lda.Opt[[1]],
         levels(test.dino.dist$Group)[1],
         levels(test.dino.dist$Group)[2]), test.dino.dist$Group
  )
knitr::kable(ldaConf.Opt)
```

| | Ornithischia | Theropoda |
|---|---|---|
| Ornithischia | 22 | 1 |
| Theropoda | 11 | 56 |

```
stats.ldaOpt <- confMatStats(ldaConf.Opt)
knitr::kable(stats.ldaOpt)
```

| | x |
|---|---|
| Sensitivity | 0.9824561 |
| Specificity | 0.6666667 |
| Accuracy | 0.8666667 |

#### Forward stepwise logistic regression (LR)

We use forward step-wise logistic regression which selects a subset of the landmarks based on the Akaike Information Criterion (AIC). Results before adjusting the threshold.

```
# Set up null model
glmNull <- glm(Group~1,data=train.dino.dist, family = "binomial")

# Define scope: Maximal set of variables: All variables
allVar <- formula(terms(Group~.,data=train.dino.dist))

fstep.glm <- step(glmNull, direction='forward', scope=allVar, trace = 0)

fstep.glm.predict<- predict(fstep.glm, test.dino.dist, type = "response")

fstep.glmConf<-table(
  ifelse(fstep.glm.predict<0.5, levels(test.dino.dist$Group)[1],
                levels(test.dino.dist$Group)[2]), test.dino.dist$Group
  )
knitr::kable(fstep.glmConf)
```

| | Ornithischia | Theropoda |
|---|---|---|
| Ornithischia | 25 | 10 |
| Theropoda | 8 | 47 |

```
stats.glm <- confMatStats(fstep.glmConf)
knitr::kable(stats.glm)
```

| | x |
|---|---|
| Sensitivity | 0.8245614 |
| Specificity | 0.7575758 |
| Accuracy | 0.8000000 |

Determining an adjusted threshold via the ROC curve.

```
par(pty="s")
glm.roc <- roc(test.dino.dist$Group, fstep.glm.predict,
               print.auc=TRUE,
               # auc.polygon=TRUE,
               plot=TRUE,
               print.auc.x=0.5,print.auc.y=0.3,
               legacy.axes=TRUE,
               xlim=c(1,0), ylim=c(0,1),
               #,main="Linear Discriminant Analysis (LDA)"
               )
glm.Opt <- coords(glm.roc, "best")
```

```r
points(glm.Opt$specificity, glm.Opt$sensitivity, col="red", pch=19, cex=1.5)
```

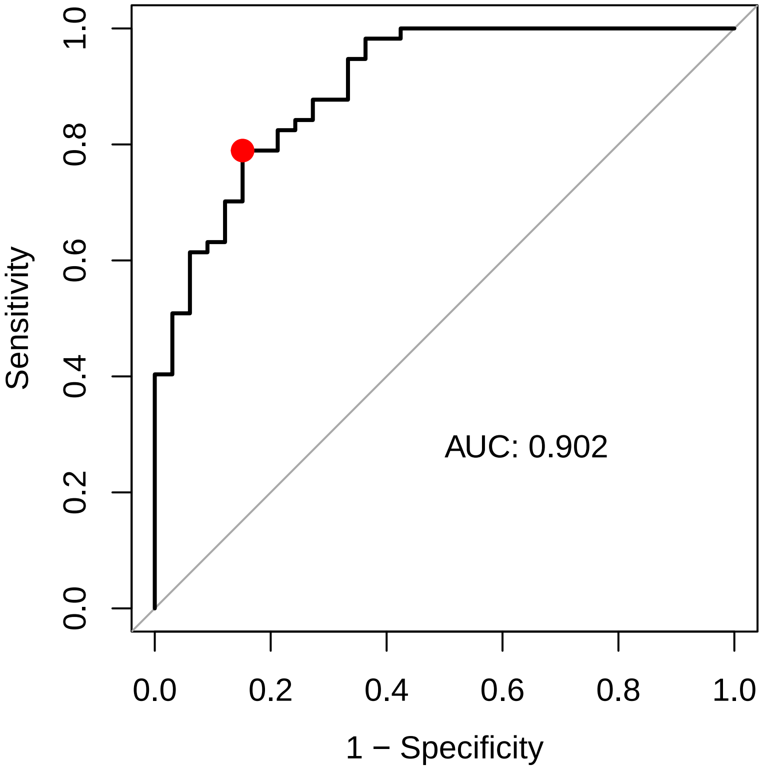


```r
if(plotPDF) {
  pdf("figures/LR_ROC.pdf")
  par(mar=c(4,4,1,1), cex=2)
  glm.forward.roc <- roc(test.dino.dist$Group, fstep.glm.predict,
                         print.auc=TRUE,
                         # auc.polygon=TRUE,
                         plot=TRUE,
                         print.auc.x=0.5,print.auc.y=0.3,
                         print.auc.cex=1,
                         legacy.axes=TRUE
               #,main="Linear Discriminant Analysis (LDA)"
              )
  points(glm.Opt$specificity, glm.Opt$sensitivity, col="red", pch=19, cex=1)
  dev.off()
}
```

Results after adjusting the threshold.

```r
glmConf.Opt<-table(
  ifelse(fstep.glm.predict<glm.Opt[[1]],
         levels(test.dino.dist$Group)[1],
         levels(test.dino.dist$Group)[2]), test.dino.dist$Group
  )
knitr::kable(glmConf.Opt)
```

| | Ornithischia | Theropoda |
|---|---|---|
| Ornithischia | 28 | 12 |
| Theropoda | 5 | 45 |

```r
stats.glmOpt <- confMatStats(glmConf.Opt)
knitr::kable(stats.glmOpt)
```

| | x |
|---|---|
| Sensitivity | 0.7894737 |
| Specificity | 0.8484848 |
| Accuracy | 0.8111111 |

Landmarks selected via stepwise logistic regression.

```r
attributes(terms(fstep.glm))$term.labels
```

```
## [1] "X20" "X2"  "X5"  "X15" "X4"  "X9"  "X10" "X17"
```

#### Multi-layer perceptron

The `caret` package is used for carrying out a grid search for hyperparameter optimisation. Results before adjusting the threshold.

```r
library("caret")

start_time <- Sys.time()
nnetKappa <- train(Group ~ ., data = train.dino.dist,
                         tuneLength=10,
                  method = "nnet",
                  metric="Kappa",
                  maxit=10000,
                  trace=FALSE)
end_time <- Sys.time()

end_time - start_time
```

```
## Time difference of 9.117483 mins
```

```r
nnetKappa.predict <- predict(nnetKappa, test.dino.dist, type="prob")
names(nnetKappa.predict$Ornithischia) <- row.names(nnetKappa.predict)
names(nnetKappa.predict$Theropoda) <- row.names(nnetKappa.predict)
nnetConf <- table(
  ifelse(nnetKappa.predict$Theropoda<0.5,
         levels(test.dino.dist$Group)[1],
         levels(test.dino.dist$Group)[2]), test.dino.dist$Group
  )
knitr::kable(nnetConf)
```

| | Ornithischia | Theropoda |
|---|---|---|
| Ornithischia | 30 | 7 |
| Theropoda | 3 | 50 |

```r
stats.mlp <- confMatStats(nnetConf)
knitr::kable(stats.mlp)
```

| | x |
|---|---|
| Sensitivity | 0.8771930 |
| Specificity | 0.9090909 |
| Accuracy | 0.8888889 |

Determining an adjusted threshold via the ROC curve.

```r
par(pty="s")
nnet.roc <- roc(test.dino.dist$Group, as.numeric(nnetKappa.predict$Theropoda),
                print.auc=TRUE,
                # auc.polygon=TRUE,
                plot=TRUE,
                print.auc.x=0.5,print.auc.y=0.3,
                legacy.axes=TRUE,
                asp=1
                #, xlim=c(1,0), ylim=c(0.6,0.4)
)
# plot(nnet.roc,
#      xlim=c(1,0), ylim=c(0,1)
# )#, main = 'Theropods vs Ornithopods')
# nnet.roc$auc

nnet.Opt <- coords(nnet.roc, "best")
# nnet.Opt

points(nnet.Opt$specificity, nnet.Opt$sensitivity, col="red", pch=19, cex=1.5)
```

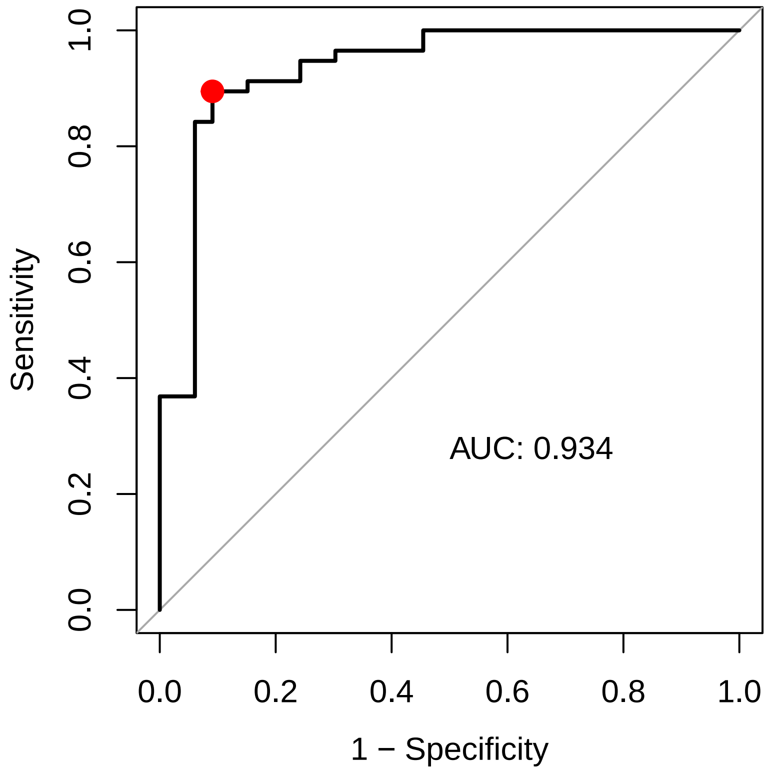


```r
if(plotPDF) {
  pdf("figures/MLP_ROC.pdf")
  par(mar=c(4,4,1,1), cex=2)
  nnet.roc <- roc(test.dino.dist$Group, as.numeric(nnetKappa.predict$Theropoda),
                print.auc=TRUE,
                # auc.polygon=TRUE,
                plot=TRUE,
                print.auc.x=0.5,print.auc.y=0.3,
                print.auc.cex=1,
```

```
                legacy.axes=TRUE,
                asp=1
                #, xlim=c(1,0), ylim=c(0.6,0.4)
  )
  points(nnet.Opt$specificity, nnet.Opt$sensitivity, col="red", pch=19, cex=1)
  dev.off()
}
```

Results after adjusting the threshold.

```
nnetConfOpt <- table(
  ifelse(nnetKappa.predict$Theropoda<nnet.Opt[[1]],
         levels(test.dino.dist$Group)[1],
         levels(test.dino.dist$Group)[2]), test.dino.dist$Group
  )
knitr::kable(nnetConfOpt)
```

| | Ornithischia | Theropoda |
|---|---|---|
| Ornithischia | 30 | 6 |
| Theropoda | 3 | 51 |

```
stats.mlpOpt <- confMatStats(nnetConfOpt)
knitr::kable(stats.mlpOpt)
```

| | x |
|---|---|
| Sensitivity | 0.8947368 |
| Specificity | 0.9090909 |
| Accuracy | 0.9000000 |

#### Random Forest

The caret package is used for carrying out a grid search for hyperparameter optimisation. Results before adjusting the threshold.

```
start_time <- Sys.time()
# Hyperparameter optimisation -> ntree?
rfMtry <- train(Group ~ ., data = train.dino.dist,
                    tuneLength=10,
                    method = "rf",
                    metric="Kappa",
                    #maxit=10000,
                    trace=FALSE)
end_time <- Sys.time()

end_time - start_time
```

```
## Time difference of 26.24083 secs
```

```
rf.predictMtry <- predict(rfMtry, test.dino.dist, type="prob")

rfConf <- table(
  ifelse(rf.predictMtry$Theropoda<0.5,
         levels(test.dino.dist$Group)[1],
         levels(test.dino.dist$Group)[2]), test.dino.dist$Group
  )
knitr::kable(rfConf)
```

| | Ornithischia | Theropoda |
|---|---|---|
| Ornithischia | 25 | 7 |
| Theropoda | 8 | 50 |

```
stats.rf <- confMatStats(rfConf)
knitr::kable(stats.rf)
```

| | x |
|---|---|
| Sensitivity | 0.8771930 |
| Specificity | 0.7575758 |
| Accuracy | 0.8333333 |

Determining an adjusted threshold via the ROC curve.

```
par(pty="s")
rf.roc <- roc(test.dino.dist$Group, rf.predictMtry$Theropoda, print.auc=TRUE,
              # auc.polygon=TRUE,
              plot=TRUE,
              print.auc.x=0.5,print.auc.y=0.3,
              legacy.axes=TRUE,
              asp=1
              #, xlim=c(1,0), ylim=c(0.6,0.4)
)
rf.Opt <- coords(rf.roc, "best")
#rf.Opt

points(rf.Opt$specificity, rf.Opt$sensitivity, col="red", pch=19, cex = 1.5)
```

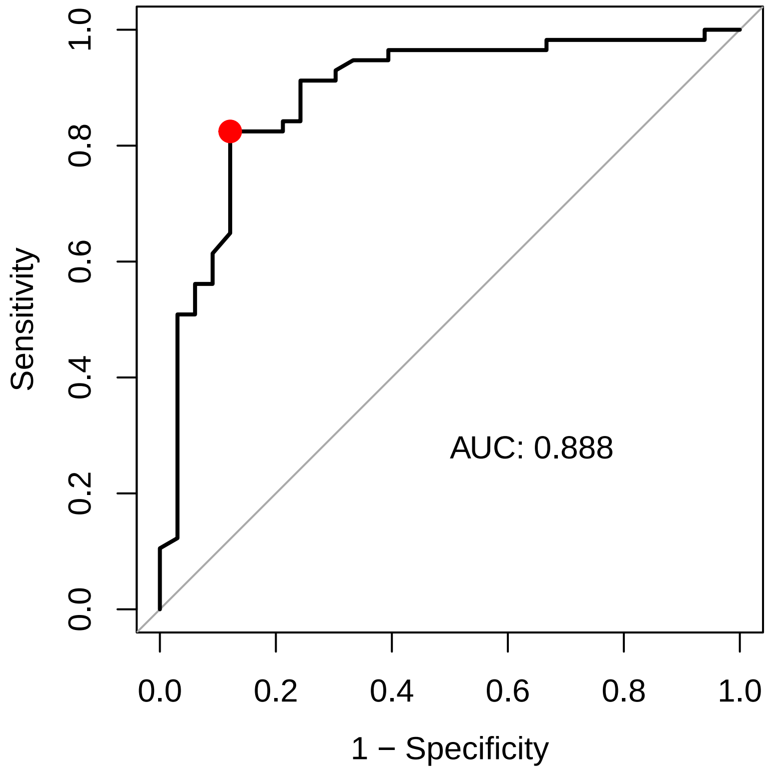


```
if(plotPDF) {
  pdf("figures/RF_ROC.pdf")
  par(mar=c(4,4,1,1), cex=2)
```

```
  rf.roc <- roc(test.dino.dist$Group, rf.predictMtry$Theropoda, print.auc=TRUE,
              # auc.polygon=TRUE,
              plot=TRUE,
              print.auc.x=0.5,print.auc.y=0.3,
              print.auc.cex=1,
              legacy.axes=TRUE,
              asp=1
              #, xlim=c(1,0), ylim=c(0.6,0.4)
    )
  points(rf.Opt$specificity, rf.Opt$sensitivity, col="red", pch=19, cex = 1)
  dev.off()
}
```

```
rfConfOpt <- table(
  ifelse(rf.predictMtry$Theropoda<rf.Opt[[1]],
         levels(test.dino.dist$Group)[1],
         levels(test.dino.dist$Group)[2]), test.dino.dist$Group)
knitr::kable(rfConfOpt)
```

| | Ornithischia | Theropoda |
|---|---|---|
| Ornithischia | 29 | 10 |
| Theropoda | 4 | 47 |

Results after adjusting the threshold.

```
stats.rfOpt <- confMatStats(rfConfOpt)
knitr::kable(stats.rfOpt)
```

| | x |
|---|---|
| Sensitivity | 0.8245614 |
| Specificity | 0.8787879 |
| Accuracy | 0.8444444 |

##### Support Vector Machine (linear)

The `caret` package is used for carrying out a grid search for hyperparameter optimisation. Results before adjusting the threshold.

```
start_time <- Sys.time()
svmKappa <- train(Group ~ ., data = train.dino.dist,
                       tuneLength=10,
                       method = "svmLinear2",
                       metric="Kappa",
                       #maxit=10000,
                      probability=TRUE,
                      decision.values=TRUE,
                       trace=FALSE)
end_time <- Sys.time()

end_time - start_time
```

```
## Time difference of 11.09548 secs
```

```
svm.predictKappa <- predict(svmKappa, test.dino.dist, type="prob")

svmConf <- table(
  ifelse(svm.predictKappa$Theropoda<0.5,
```

```
         levels(test.dino.dist$Group)[1],
         levels(test.dino.dist$Group)[2]),
  test.dino.dist$Group
  )
knitr::kable(svmConf)
```

| | Ornithischia | Theropoda |
|---|---|---|
| Ornithischia | 24 | 12 |
| Theropoda | 9 | 45 |

```
knitr::kable(confMatStats(svmConf))
```

| | x |
|---|---|
| Sensitivity | 0.7894737 |
| Specificity | 0.7272727 |
| Accuracy | 0.7666667 |

Determining an adjusted threshold via the ROC curve.

```
par( pty="s")
svm.roc <- roc(test.dino.dist$Group, as.numeric(svm.predictKappa$Theropoda),
               print.auc=TRUE,
               # auc.polygon=TRUE,
               plot=TRUE,
               print.auc.x=0.5,print.auc.y=0.3,
               legacy.axes=TRUE,
               asp=1
               #, xlim=c(1,0), ylim=c(0.6,0.4)
)

svm.Opt <- coords(svm.roc, "best")

points(svm.Opt$specificity, svm.Opt$sensitivity, col="red", pch=19, cex = 1.5)
```

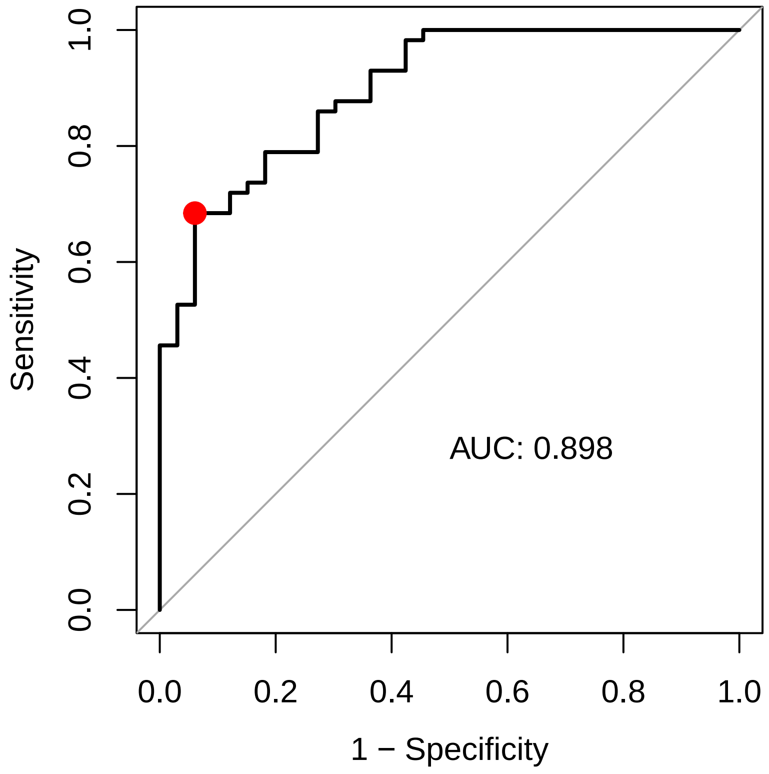


```
if(plotPDF) {
  pdf("figures/SVMlinear_ROC.pdf")
  par(mar=c(4,4,1,1), cex=2)
  svm.roc <- roc(test.dino.dist$Group, as.numeric(svm.predictKappa$Theropoda),
                 print.auc=TRUE,
                 # auc.polygon=TRUE,
                 plot=TRUE,
                 print.auc.x=0.5,print.auc.y=0.3,
                 print.auc.cex=1,
                 legacy.axes=TRUE,
                 asp=1
                 #, xlim=c(1,0), ylim=c(0.6,0.4)
    )
  points(svm.Opt$specificity, svm.Opt$sensitivity, col="red", pch=19, cex = 1)
  dev.off()
}
```

Results after adjusting the threshold.

```
svmConfOpt <- table(
  ifelse(svm.predictKappa$Theropoda<svm.Opt$threshold,
         levels(test.dino.dist$Group)[1],
        levels(test.dino.dist$Group)[2]), test.dino.dist$Group
  )
knitr::kable(svmConfOpt)
```

| | Ornithischia | Theropoda |
|---|---:|---:|
| Ornithischia | 31 | 18 |
| Theropoda | 2 | 39 |

```r
knitr::kable(confMatStats(svmConfOpt))
```

| | x |
|---|---|
| Sensitivity | 0.6842105 |
| Specificity | 0.9393939 |
| Accuracy | 0.7777778 |

**Support Vector Machine (radial kernel)**

The caret package is used for carrying out a grid search for hyperparameter optimisation. Results before adjusting the threshold.

```r
start_time <- Sys.time()
svmRadialKappa <- train(Group ~ ., data = train.dino.dist,
                        tuneLength=10,
                        method = "svmRadialSigma",
                        metric="Kappa",
                        #maxit=10000,
                       prob.model=TRUE,
                       decision.values=TRUE,
                        trace=FALSE)
end_time <- Sys.time()

end_time - start_time
```

```
## Time difference of 46.16281 secs
```

```r
svmRadial.predictKappa <- predict(svmRadialKappa, test.dino.dist, type="prob")

svmRadialConf <- table(ifelse(svmRadial.predictKappa$Theropoda<0.5,
                              levels(test.dino.dist$Group)[1],
                              levels(test.dino.dist$Group)[2]),
                       test.dino.dist$Group)
knitr::kable(svmRadialConf)
```

| | Ornithischia | Theropoda |
|---|---|---|
| Ornithischia | 26 | 7 |
| Theropoda | 7 | 50 |

```r
stats.svm <- confMatStats(svmRadialConf)
knitr::kable(stats.svm)
```

| | x |
|---|---|
| Sensitivity | 0.8771930 |
| Specificity | 0.7878788 |
| Accuracy | 0.8444444 |

Determining an adjusted threshold via the ROC curve.

```r
par(pty="s")
svmRadial.roc <- roc(test.dino.dist$Group, svmRadial.predictKappa$Theropoda,
               print.auc=TRUE,
               # auc.polygon=TRUE,
               plot=TRUE,
               print.auc.x=0.5,print.auc.y=0.3,
               legacy.axes=TRUE,
               asp=1
               #, xlim=c(1,0), ylim=c(0.6,0.4)
```

```
)

svmRadial.Opt <- coords(svmRadial.roc, "best")

points(svmRadial.Opt$specificity, svmRadial.Opt$sensitivity,
       col="red", pch=19, cex = 1.5)
```

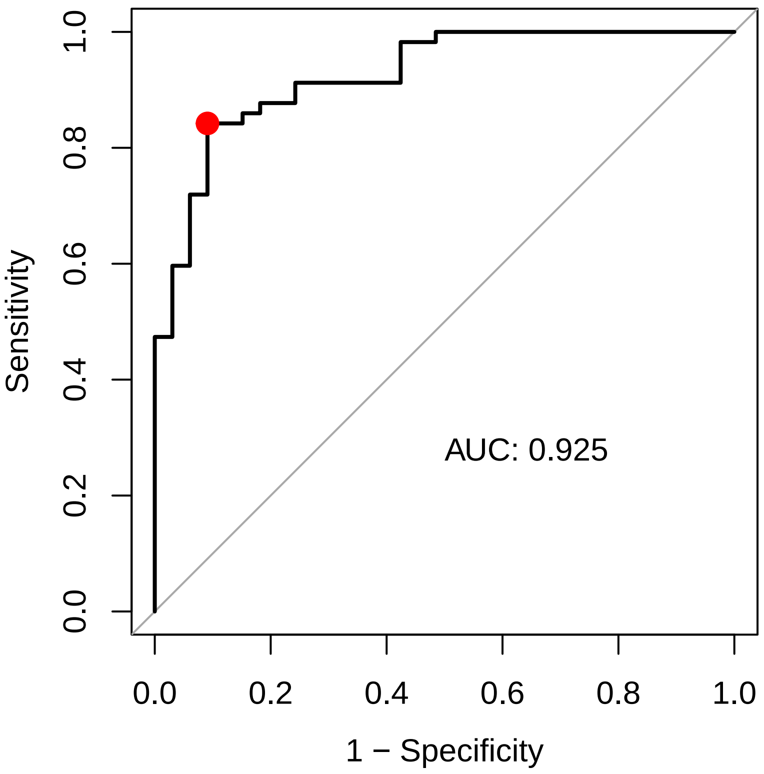


```
if(plotPDF) {
  pdf("figures/SVMradSigma_ROC.pdf")
  par(mar=c(4,4,1,1), cex=2)
  svmRadial.roc <- roc(test.dino.dist$Group, svmRadial.predictKappa$Theropoda,
                  print.auc=TRUE,
                  # auc.polygon=TRUE,
                  plot=TRUE,
                  print.auc.x=0.5,print.auc.y=0.3,
                  print.auc.cex = 1,
                  legacy.axes=TRUE,
                  asp=1
                  #, xlim=c(1,0), ylim=c(0.6,0.4)
  )
  points(svmRadial.Opt$specificity, svmRadial.Opt$sensitivity,
         col="red", pch=19, cex = 1)
  dev.off()
}
```

Results after adjusting the threshold.

```
svmRadialConfOpt <- table(
  ifelse(svmRadial.predictKappa$Theropoda<svm.Opt$threshold,
         levels(test.dino.dist$Group)[1],
```

```
        levels(test.dino.dist$Group)[2]), test.dino.dist$Group
  )
knitr::kable(svmRadialConfOpt)
```

| | Ornithischia | Theropoda |
|---|---|---|
| Ornithischia | 28 | 9 |
| Theropoda | 5 | 48 |

```
stats.svmOpt <- confMatStats(svmRadialConfOpt)
knitr::kable(stats.svmOpt)
```

| | x |
|---|---|
| Sensitivity | 0.8421053 |
| Specificity | 0.8484848 |
| Accuracy | 0.8444444 |

#### MARS

Results before adjusting the threshold.

```
library(mda)
marsResults <- fda(Group~., data = train.dino.dist, method = mars)

# Obtain probabilities -> Second columns contains probability for Theropods
marsScores <- predict(marsResults, test.dino.dist,type = "posterior")
#marsScores[,2]
marsConf <- table(ifelse(marsScores[,2]<0.5, levels(test.dino.dist$Group)[1],
                    levels(test.dino.dist$Group)[2]), test.dino.dist$Group)
knitr::kable(marsConf)
```

| | Ornithischia | Theropoda |
|---|---|---|
| Ornithischia | 26 | 10 |
| Theropoda | 7 | 47 |

```
stats.mars <- confMatStats(marsConf)
knitr::kable(stats.mars)
```

| | x |
|---|---|
| Sensitivity | 0.8245614 |
| Specificity | 0.7878788 |
| Accuracy | 0.8111111 |

Determining an adjusted threshold via the ROC curve.

```
par(pty="s")
mars.roc <- roc(test.dino.dist$Group, marsScores[,2],
                print.auc=TRUE,
                # auc.polygon=TRUE,
                plot=TRUE,
                print.auc.x=0.5,print.auc.y=0.3,
                legacy.axes=TRUE,
                asp=1
                #,main="Linear Discriminant Analysis (LDA)"
)

mars.Opt <- coords(mars.roc, "best")
#mars.Opt
```

```
points(mars.Opt$specificity, mars.Opt$sensitivity, col="red", pch=19, cex = 1.5)
```

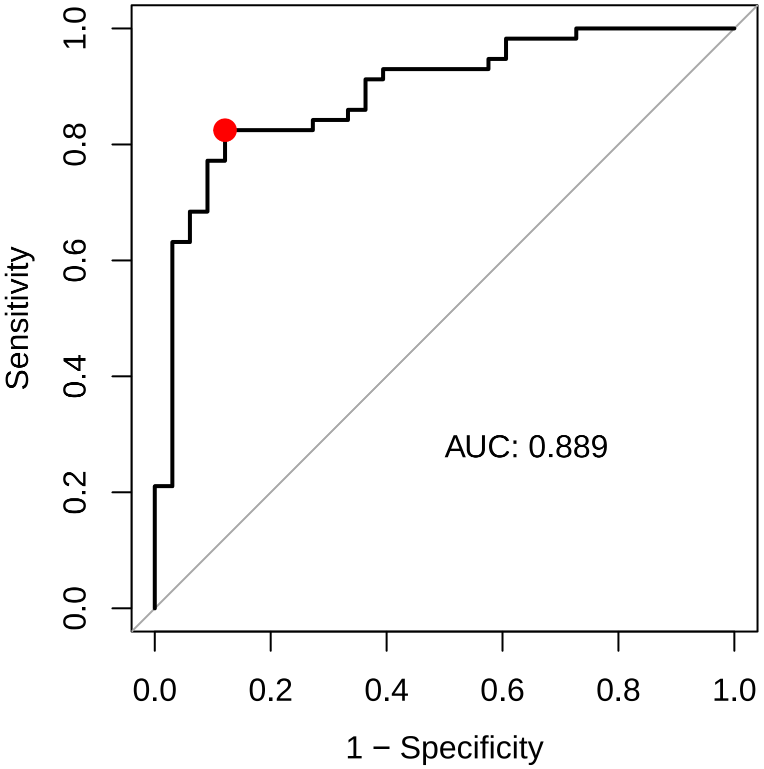


```
if(plotPDF) {
  pdf("figures/MARS_ROC.pdf")
  par(mar=c(4,4,1,1), cex=2)
  mars.roc <- roc(test.dino.dist$Group, marsScores[,2],
                print.auc=TRUE,
                # auc.polygon=TRUE,
                plot=TRUE,
                print.auc.x=0.5,print.auc.y=0.3,
                print.auc.cex=1,
                legacy.axes=TRUE,
                asp=1
                #,main="Linear Discriminant Analysis (LDA)"
  )
  points(mars.Opt$specificity, mars.Opt$sensitivity, col="red", pch=19, cex = 1)
  dev.off()
}
```

Results after adjusting the threshold.

```
marsConfOpt <- table(
  ifelse(marsScores[,2]<mars.Opt$threshold, levels(test.dino.dist$Group)[1],
                  levels(test.dino.dist$Group)[2]), test.dino.dist$Group)
knitr::kable(marsConfOpt)
```

| | Ornithischia | Theropoda |
|---|---|---|
| Ornithischia | 29 | 10 |
| Theropoda | 4 | 47 |

```r
stats.marsOpt <- confMatStats(marsConfOpt)
knitr::kable(stats.marsOpt)
```

| | x |
|---|---|
| Sensitivity | 0.8245614 |
| Specificity | 0.8787879 |
| Accuracy | 0.8444444 |

#### Plots of performance metrics

```r
metrics <- data.frame(
  rbind(
    stats.glm, stats.glmOpt,
    stats.mlp, stats.mlpOpt,
    stats.lda, stats.ldaOpt,
    stats.svm, stats.svmOpt,
    stats.rf, stats.rfOpt,
    stats.mars, stats.marsOpt
  )
)

# layout(matrix(1:6, nrow = 3, ncol = 2, byrow = TRUE))
# for (i in 1:6) {
#   #par(pty="s")
#   matplot(metrics[c(2*i-1,2*i), ], xlim=c(0.5,2.5), ylim = c(0.65,1),
#           pch = 1, type = "b", col = c("black","red",  "blue"), lty=c(2,2,1),
#           ylab="", main=rownames(metrics[2*i-1, ]), xaxt="n"#, asp=1
#           )
#     axis(side=1,at=c(1,2),labels=c(expression(paste(theta,"=0.5")), expression(paste(theta, " adjuste
# }


par(mfrow=c(1,2))
for (i in 1:6) {
  #par(pty="s")
  matplot(metrics[c(2*i-1,2*i), ], xlim=c(0.5,2.5), ylim = c(0.65,1),
          pch = 1, type = "b", col = c("black","red",  "blue"), lty=c(2,2,1),
          ylab="", main=rownames(metrics[2*i-1, ]), xaxt="n", asp=1
          )
    axis(side=1,at=c(1,2),labels=c(expression(paste(theta,"=0.5")),
                                   expression(paste(theta, " adjusted"))))
}
```

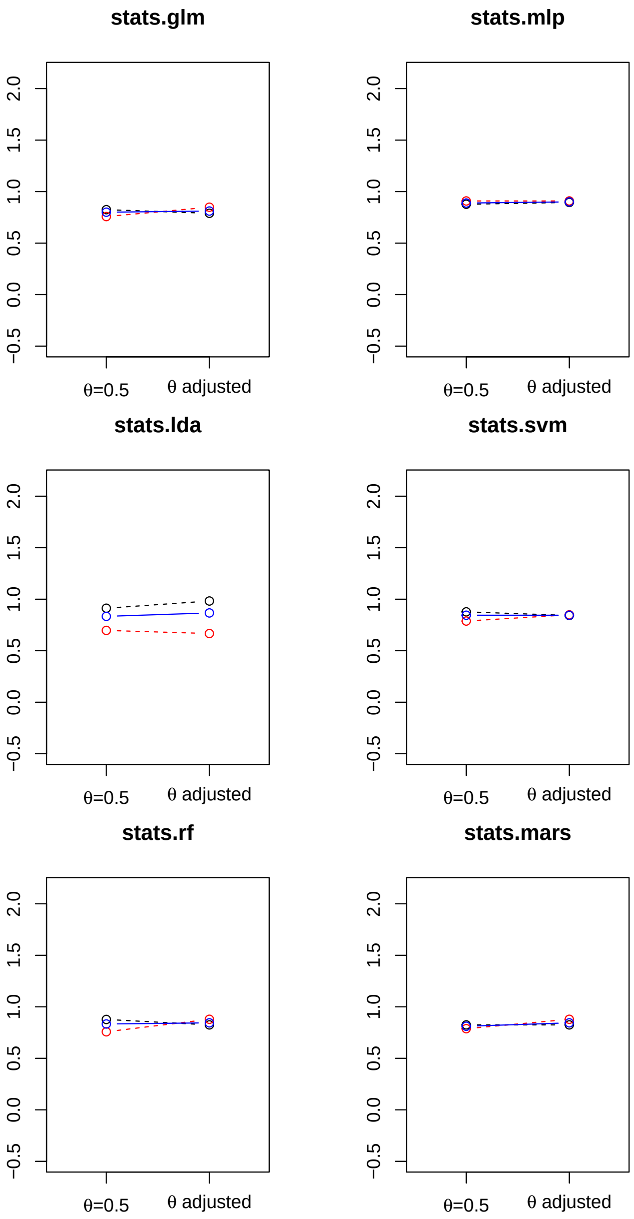


```
if(plotPDF) {
  for (i in 1:6) {
    pdf(sprintf("figures/%s.pdf", rownames(metrics[2*i-1, ])))
    par(mar=c(4,4,1,1), cex=2)
    matplot(metrics[c(2*i-1,2*i), ], xlim=c(0.5,2.5), ylim = c(0.65,1),
            pch = 1, type = "b", col = c("black","red",  "blue"), lty=c(2,2,1),
            ylab="", xaxt="n")
    axis(side=1,at=c(1,2),labels=c(expression(paste(theta,"=0.5")), expression(paste(theta, " adjusted"
    dev.off()
  }
}
```

### Misclassified samples

For each method, the samples that are misclassified are determined.

**Linear Discrimiant Analysis**

```
lda.false.orn <- which(
  (lda.predict$posterior[,2]>0.5) &
    (test.dino.dist$Group==levels(test.dino.dist$Group)[1])
                       )
lda.false.orn
```

```
##  12  13  20  28  30  65  89 154 155 213
##   5   6   7  10  11  22  28  49  50  64
```

```
lda.false.ther <- which(
  (lda.predict$posterior[,2]<0.5) &
    (test.dino.dist$Group==levels(test.dino.dist$Group)[2]))
lda.false.ther
```

```
## 147 192 222 251 273
##  47  60  67  79  83
```

```
# Summarise indices of all misclassified samples
lda.misclass <- as.integer(names(c(lda.false.orn, lda.false.ther)))

# Misclassified samples
lda.false.orn.opt <- which(
  (lda.predict$posterior[,2]>lda.Opt$threshold) &
    (test.dino.dist$Group==levels(test.dino.dist$Group)[1])
  )
lda.false.orn.opt
```

```
##  12  13  20  28  30  65  89 154 155 213 290
##   5   6   7  10  11  22  28  49  50  64  89
```

```
lda.false.ther.opt <- which(
  (lda.predict$posterior[,2]<lda.Opt$threshold) &
    (test.dino.dist$Group==levels(test.dino.dist$Group)[2])
  )
lda.false.ther.opt
```

```
## 192
##  60
```

```
# Summarise indices of all misclassified samples
lda.misclass.opt <- as.integer(names(c(lda.false.orn.opt, lda.false.ther.opt)))
```

**Forward stepwise regression**

```
# Misclassified ornithopods
glm.false.orn <- which(
  (fstep.glm.predict>0.5) &
    (test.dino.dist$Group==levels(test.dino.dist$Group)[1])
  )
glm.false.orn
```

```
##  12  13  28  30  65  89 154 213
##   5   6  10  11  22  28  49  64
```

```
# Misclassified theropods
glm.false.ther <-which(
```

```
  (fstep.glm.predict<0.5) &
    (test.dino.dist$Group==levels(test.dino.dist$Group)[2])
  )
glm.false.ther
```

```
##  79  96  99 147 149 203 222 251 273 296
##  25  32  33  47  48  62  67  79  83  90
```

```
# Summarise indices of all misclassified samples
glm.misclass <- as.integer(names(c(glm.false.orn, glm.false.ther)))

# Misclassified ornithopods
glm.false.orn.opt <- which(
  (fstep.glm.predict>glm.Opt$threshold) &
    (test.dino.dist$Group==levels(test.dino.dist$Group)[1])
  )
glm.false.orn.opt
```

```
##  12  13  65  89 154
##   5   6  22  28  49
```

```
# Misclassified theropods
glm.false.ther.opt <-which(
  (fstep.glm.predict<glm.Opt$threshold) &
    (test.dino.dist$Group==levels(test.dino.dist$Group)[2])
  )
glm.false.ther.opt
```

```
##  64  79  96  99 147 149 203 222 251 273 276 296
##  21  25  32  33  47  48  62  67  79  83  84  90
```

```
# Summarise indices of all misclassified samples
glm.misclass.opt <- as.integer(names(c(glm.false.orn.opt, glm.false.ther.opt)))
glm.misclass.opt
```

```
##  [1]  12  13  65  89 154  64  79  96  99 147 149 203 222 251 273 276 296
```

#### Multi-layer perceptron

```
# Misclassified ornithopods -> Slightly silly way of using setNames... changing directly in data frame
nnet.false.orn <- which(
  (setNames(nnetKappa.predict$Ornithischia,row.names(nnetKappa.predict))<0.5) &
    (test.dino.dist$Group==levels(test.dino.dist$Group)[1])
  )
nnet.false.orn
```

```
## 12 20 89
##  5  7 28
```

```
# Misclassified theropods
nnet.false.ther <-which(
  (setNames(nnetKappa.predict$Theropoda,row.names(nnetKappa.predict))<0.5) &
    (test.dino.dist$Group==levels(test.dino.dist$Group)[2])
  )
nnet.false.ther
```

```
##  80  96  99 128 251 273 276
```

```
##  26  32  33  40  79  83  84
```

```
# Summarise indices of all misclassified samples
nnet.misclass <- as.integer(names(c(nnet.false.orn, nnet.false.ther)))
nnet.misclass
```

```
##  [1]  12  20  89  80  96  99 128 251 273 276
```

```
# Misclassified ornithopods -> Slightly silly way of using setNames... changing directly in data frame
nnet.false.orn.opt <- which(
  (setNames(nnetKappa.predict$Ornithischia,row.names(nnetKappa.predict))<nnet.Opt$threshold) &
    (test.dino.dist$Group==levels(test.dino.dist$Group)[1])
  )
nnet.false.orn.opt
```

```
## 12 20 89
##  5  7 28
```

```
# Misclassified theropods
nnet.false.ther.opt <-which(
  (setNames(nnetKappa.predict$Theropoda,row.names(nnetKappa.predict))<nnet.Opt$threshold) &
    (test.dino.dist$Group==levels(test.dino.dist$Group)[2])
                            )
nnet.false.ther.opt
```

```
##  80  96  99 251 273 276
##  26  32  33  79  83  84
```

```
# Summarise indices of all misclassified samples
nnet.misclass.opt <- as.integer(names(c(nnet.false.orn.opt, nnet.false.ther.opt)))
nnet.misclass.opt
```

```
## [1]  12  20  89  80  96  99 251 273 276
```

#### Random Forest

```
# Misclassified ornithopods
rf.false.orn <- which(
  (setNames(rf.predictMtry$Ornithischia<0.5, row.names(rf.predictMtry))) &
    (test.dino.dist$Group==levels(test.dino.dist$Group)[1])
  )
rf.false.orn
```

```
##  12  13  20  89 154 173 213 290
##   5   6   7  28  49  55  64  89
```

```
# Misclassified theropods
rf.false.ther <-which(
  (setNames(rf.predictMtry$Theropoda<0.5, row.names(rf.predictMtry))) &
    (test.dino.dist$Group==levels(test.dino.dist$Group)[2])
  )
rf.false.ther
```

```
##  80  96 121 146 167 273 276
##  26  32  37  46  52  83  84
```

```
# Summarise indices of all misclassified samples
rf.misclass <- as.integer(names(c(rf.false.orn, rf.false.ther)))
rf.misclass
```

```
##  [1]  12  13  20  89 154 173 213 290  80  96 121 146 167 273 276
```

```
# Misclassified ornithopods
rf.false.orn.opt <- which(
  (setNames(rf.predictMtry$Ornithischia<rf.Opt$threshold, row.names(rf.predictMtry))) &
    (test.dino.dist$Group==levels(test.dino.dist$Group)[1])
  )
rf.false.orn.opt
```

```
##  12  13  20  89 134 154 173 213 290
##   5   6   7  28  43  49  55  64  89
```

```
# Misclassified theropods
rf.false.ther.opt <-which(
  (setNames(rf.predictMtry$Theropoda<rf.Opt$threshold, row.names(rf.predictMtry))) &
    (test.dino.dist$Group==levels(test.dino.dist$Group)[2])
  )
rf.false.ther.opt
```

```
##  50  80  96  99 121 146 167 251 273 276
##  17  26  32  33  37  46  52  79  83  84
```

```
# Summarise indices of all misclassified samples
rf.misclass.opt <- as.integer(names(c(rf.false.orn.opt, rf.false.ther.opt)))
rf.misclass.opt
```

```
##  [1]  12  13  20  89 134 154 173 213 290  50  80  96  99 121 146 167 251 273 276
```

**Support Vector Machine**

```
# Misclassified ornithopods
# We need to use rf.predictMtry to find the indices, svm seems to destroy these indices!
svm.false.orn <- which(
  (setNames(svm.predictKappa$Theropoda>0.5, row.names(rf.predictMtry))) &
    (test.dino.dist$Group==levels(test.dino.dist$Group)[1])
  )
svm.false.orn
```

```
##  12  13  28  30  65  89 154 155 213
##   5   6  10  11  22  28  49  50  64
```

```
# Misclassified theropods
svm.false.ther <-which(
  (setNames(svm.predictKappa$Theropoda<0.5, row.names(rf.predictMtry))) &
    (test.dino.dist$Group==levels(test.dino.dist$Group)[2])
  )
svm.false.ther
```

```
##  79  80  96  99 147 149 203 222 251 273 276 296
##  25  26  32  33  47  48  62  67  79  83  84  90
```

```
svm.misclass <- as.integer(names(c(svm.false.orn, svm.false.ther)))
svm.misclass
```

```
##  [1]  12  13  28  30  65  89 154 155 213  79  80  96  99 147 149 203 222 251 273
## [20] 276 296
```

```
# Misclassified ornithopods
# We need to use rf.predictMtry to find the indices, svm seems to destroy these indices!
```

```
svm.false.orn.opt <- which(
  (setNames(svm.predictKappa$Theropoda>svm.Opt$threshold, row.names(rf.predictMtry))) &
    (test.dino.dist$Group==levels(test.dino.dist$Group)[1])
  )
svm.false.orn.opt
```

```
## 12 89
##  5 28
```

```
# Misclassified theropods
svm.false.ther.opt <-which(
  (setNames(svm.predictKappa$Theropoda<svm.Opt$threshold, row.names(rf.predictMtry))) &
    (test.dino.dist$Group==levels(test.dino.dist$Group)[2])
  )
svm.false.ther.opt
```

```
##  41  47  64  79  80  96  99 128 147 149 192 203 222 251 266 273 276 296
##  14  16  21  25  26  32  33  40  47  48  60  62  67  79  80  83  84  90
```

```
svm.misclass.opt <- as.integer(names(c(svm.false.orn.opt, svm.false.ther.opt)))
svm.misclass.opt
```

```
##  [1]  12  89  41  47  64  79  80  96  99 128 147 149 192 203 222 251 266 273 276
## [20] 296
```

#### MARS

```
# Misclassified ornithopods
# We need to use rf.predictMtry to get the indices!
mars.false.orn <- which(
  (setNames(marsScores[,2]>0.5, row.names(rf.predictMtry))) &
    (test.dino.dist$Group==levels(test.dino.dist$Group)[1])
  )
mars.false.orn
```

```
##  12  13  20  27  89 154 213
##   5   6   7   9  28  49  64
```

```
# Misclassified theropods
mars.false.ther <-which(
  (setNames(marsScores[,2]<0.5, row.names(rf.predictMtry))) &
    (test.dino.dist$Group==levels(test.dino.dist$Group)[2])
  )
mars.false.ther
```

```
##   4  80  96  99 121 221 222 251 273 276
##   2  26  32  33  37  66  67  79  83  84
```

```
mars.misclass <- as.integer(names(c(mars.false.orn, mars.false.ther)))
mars.misclass
```

```
##  [1]  12  13  20  27  89 154 213   4  80  96  99 121 221 222 251 273 276
```

```
# Misclassified ornithopods
# We need to use rf.predictMtry to get the indices!
mars.false.orn.opt <- which(
  (setNames(marsScores[,2]>mars.Opt$threshold, row.names(rf.predictMtry))) &
    (test.dino.dist$Group==levels(test.dino.dist$Group)[1])
```

```r
  )
mars.false.orn.opt
```

```
##  12  20  89 154
##   5   7  28  49
```

```r
# Misclassified theropods
mars.false.ther.opt <-which(
  (setNames(marsScores[,2]<mars.Opt$threshold, row.names(rf.predictMtry))) &
    (test.dino.dist$Group==levels(test.dino.dist$Group)[2])
  )
mars.false.ther.opt
```

```
##   4  80  96  99 121 221 222 251 273 276
##   2  26  32  33  37  66  67  79  83  84
```

```r
mars.misclass.opt <- as.integer(names(c(mars.false.orn.opt, mars.false.ther.opt)))
mars.misclass.opt
```

```
##  [1]  12  20  89 154   4  80  96  99 121 221 222 251 273 276
```

#### Summarise all misclassified samples

Misclassied samples for all methods are summarised in a table. We determine which footprints have been misclassified by all six methods…

```r
classifiers <- read.csv("classifiers.csv", header=T, row.names=1)
classifiers <- classifiers[-which(is.na(classifiers[,4])), ]

all.misclass <- sort(unique(c(lda.misclass, glm.misclass, nnet.misclass,
                              rf.misclass, svm.misclass, mars.misclass)))
all.misclass
```

```
##  [1]   4  12  13  20  27  28  30  65  79  80  89  96  99 121 128 146 147 149 154
## [20] 155 167 173 192 203 213 221 222 251 273 276 290 296
```

```r
misclassTable <- classifiers[all.misclass, ]
misclassTable$lda <- (all.misclass %in% lda.misclass)
misclassTable$glm <- (all.misclass %in% glm.misclass)
misclassTable$nnet <- (all.misclass %in% nnet.misclass)
misclassTable$rf <- (all.misclass %in% rf.misclass)
misclassTable$svm <- (all.misclass %in% svm.misclass)
misclassTable$mars <- (all.misclass %in% mars.misclass)
misclassTable$indices <- all.misclass

# Quantify which samples are frequently misclassified
misclassTable$nMisclass <- rowSums(misclassTable[,c("lda", "glm", "nnet", "rf", "svm", "mars")])

# Drop irrelevant columns for output
misclassTableOut <- subset(misclassTable, select = -c(SizeBin, ProjBin))

knitr::kable(misclassTableOut) %>%   kable_styling(latex_options = "scale_down")
```

### Plots of misclassified footprints

Footprints that were misclassified by more than half of the methods will be visualised using wireframe graphs.

| | Ichnogenus | Period | Epoch | Group | lda | glm | nnet | rf | svm | mars | indices | nMisclass |
|---|---|---|---|---|---|---|---|---|---|---|---|---|
| barco 2005 1 | Iberosauripus | Cretaceous | Cretaceous Early | Theropoda | FALSE | FALSE | FALSE | FALSE | FALSE | TRUE | 4 | 1 |
| castanera 2013b 1 | NA | NA | NA | Ornithischia | TRUE | TRUE | TRUE | TRUE | TRUE | TRUE | 12 | 6 |
| castanera 2013b 2 | NA | NA | NA | Ornithischia | TRUE | TRUE | FALSE | TRUE | TRUE | TRUE | 13 | 5 |
| dalman 2013 1 | Anomoepus | Jurassic | Jurassic Early | Ornithischia | TRUE | FALSE | TRUE | TRUE | FALSE | TRUE | 20 | 4 |
| ellenberger 1974 2 | Moyenisauropus | Jurassic | Jurassic Early | Ornithischia | FALSE | FALSE | FALSE | FALSE | FALSE | TRUE | 27 | 1 |
| ellenberger 1974 3 | Moyenisauropus | Jurassic | Jurassic Early | Ornithischia | TRUE | TRUE | FALSE | FALSE | TRUE | FALSE | 28 | 3 |
| ellenberger 1974 5 | Moyenisauropus | Jurassic | Jurassic Early | Ornithischia | TRUE | TRUE | FALSE | FALSE | TRUE | FALSE | 30 | 3 |
| gierlinski 2004 6 | Anomoepus | Jurassic | Jurassic Ealry | Ornithischia | TRUE | TRUE | FALSE | FALSE | TRUE | FALSE | 65 | 3 |
| gierlinski 2009b 3 | Therangospodus | Jurassic | Jurassic Middle | Theropoda | FALSE | TRUE | FALSE | FALSE | TRUE | FALSE | 79 | 2 |
| gierlinski 2009b 4 | Carmelopodus | Jurassic | Jurassic Middle | Theropoda | FALSE | FALSE | TRUE | TRUE | TRUE | TRUE | 80 | 4 |
| jianjun 2012 3 | Anomoepus | Jurassic | NA | Ornithischia | TRUE | TRUE | TRUE | TRUE | TRUE | TRUE | 89 | 6 |
| kim 2017 3 | Corpulentatpus | Cretaceous | Cretaceous Early | Theropoda | FALSE | TRUE | TRUE | TRUE | TRUE | TRUE | 96 | 5 |
| lallensack 2016 2 | NA | Cretaceous | Cretaceous Early | Theropoda | FALSE | TRUE | TRUE | FALSE | TRUE | TRUE | 99 | 4 |
| lockley 1998b 2 | Megalosauripus | Jurassic | Jurassic Late | Theropoda | FALSE | FALSE | FALSE | TRUE | FALSE | TRUE | 121 | 2 |
| lockley 1998c 5 | Therangospodus | Jurassic | Jurassic Late | Theropoda | FALSE | FALSE | TRUE | FALSE | FALSE | FALSE | 128 | 1 |
| lockley 2008 4 | Hispanosauropus | Jurassic | Jurassic Late | Theropoda | FALSE | FALSE | FALSE | TRUE | FALSE | FALSE | 146 | 1 |
| lockley 2008b 1 | Minisauripus | Cretaceous | Cretaceous Late | Theropoda | TRUE | TRUE | FALSE | FALSE | TRUE | FALSE | 147 | 3 |
| lockley 2008b 3 | Minisauripus | Cretaceous | Cretaceous Early | Theropoda | FALSE | TRUE | FALSE | FALSE | TRUE | FALSE | 149 | 2 |
| lockley 2009 4 | Dinehichnus | Jurassic | Jurassic Early | Ornithischia | TRUE | TRUE | FALSE | TRUE | TRUE | TRUE | 154 | 5 |
| lockley 2009 5 | Dinehichnus | Jurassic | Jurassic Early | Ornithischia | TRUE | FALSE | FALSE | FALSE | TRUE | FALSE | 155 | 2 |
| lockley 2014b 1 | Irenesauripus | Cretaceous | NA | Theropoda | FALSE | FALSE | FALSE | TRUE | FALSE | FALSE | 167 | 1 |
| lockley 2014e 1 | NA | Cretaceous | NA | Ornithischia | FALSE | FALSE | FALSE | TRUE | FALSE | FALSE | 173 | 1 |
| matsukawa 2006 6 | NA | Cretaceous | Cretaceous Early | Theropoda | TRUE | FALSE | FALSE | FALSE | FALSE | FALSE | 192 | 1 |
| olsen 1980 2 | Grallator | Jurassic | Jurassic Early | Theropoda | FALSE | TRUE | FALSE | FALSE | TRUE | FALSE | 203 | 2 |
| olsen 2003 3 | Anomoepus | Jurassic | Jurassic Early | Ornithischia | TRUE | TRUE | FALSE | TRUE | TRUE | TRUE | 213 | 5 |
| pittman 1989 3 | NA | Cretaceous | Cretaceous Early | Theropoda | FALSE | FALSE | FALSE | FALSE | FALSE | TRUE | 221 | 1 |
| raath 1972 1 | NA | NA | NA | Theropoda | TRUE | TRUE | FALSE | FALSE | TRUE | TRUE | 222 | 4 |
| xing 2011 4 | Kayentapus | Cretaceous | Cretaceous Early | Theropoda | TRUE | TRUE | TRUE | FALSE | TRUE | TRUE | 251 | 5 |
| xing 2014f 1 | Paracorpulentapus | Cretaceous | Cretaceous Late | Theropoda | TRUE | TRUE | TRUE | TRUE | TRUE | TRUE | 273 | 6 |
| xing 2014h 1 | NA | Jurassic | Jurassic Early | Theropoda | FALSE | FALSE | TRUE | TRUE | TRUE | TRUE | 276 | 4 |
| xing 2016c 1 | Anomoepus | Jurassic | Jurassic Early | Ornithischia | FALSE | FALSE | FALSE | TRUE | FALSE | FALSE | 290 | 1 |
| xing 2016g 1 | Minisauripus | Cretaceous | Cretaceous Early | Theropoda | FALSE | TRUE | FALSE | FALSE | TRUE | FALSE | 296 | 2 |

```r
# misclassTable[misclassTable$nMisclass==6,]
lmArc1 <- c(4, 7, 6, 5, 1, 8, 9)
lmArc2 <- c(20, 19, 18, 2, 15, 16, 17)
lmArc3 <- c(14, 13, 3, 10, 11, 12,4)

# TODO: Code as function!
nTracks <- nrow(misclassTable[misclassTable$nMisclass==6,])

nTrackCols <- 3

lenTrackVector <- ceiling(nTracks/nTrackCols)
trackVector <- integer(lenTrackVector)

trackVector[1:nTracks] <- 1:nTracks

misclassIndicesGroup <- subset(misclassTable,
                               subset = nMisclass==6,
                               select = c(Group, indices))


layout(
  matrix(
    trackVector, ncol = 3, byrow = TRUE)
)
par(pty="s")
for (i in 1:nTracks ) {
  footprint <- subset(
      dinoprints, select = c(X, Y),
      subset = id==misclassIndicesGroup[i,"indices"]
      )
  plot(footprint,
```

```
    col=ifelse(misclassIndicesGroup[i,"Group"]=="Theropoda", "black", "red"),
    main=rownames(misclassIndicesGroup)[i]#=misclassIndicesGroup[i,"indices"],
    ,pch = 19, asp =1
    )

    lines(footprint[lmArc1, ],
          type="b",
          col=ifelse(misclassIndicesGroup[i,"Group"]=="Theropoda",
                     "black", "red"))

    lines(footprint[lmArc2, ],
          type="b",
          col=ifelse(misclassIndicesGroup[i,"Group"]=="Theropoda",
                     "black", "red"))

    lines(footprint[lmArc3, ],
          type="b",
          col=ifelse(misclassIndicesGroup[i,"Group"]=="Theropoda",
                     "black", "red"))

}
```

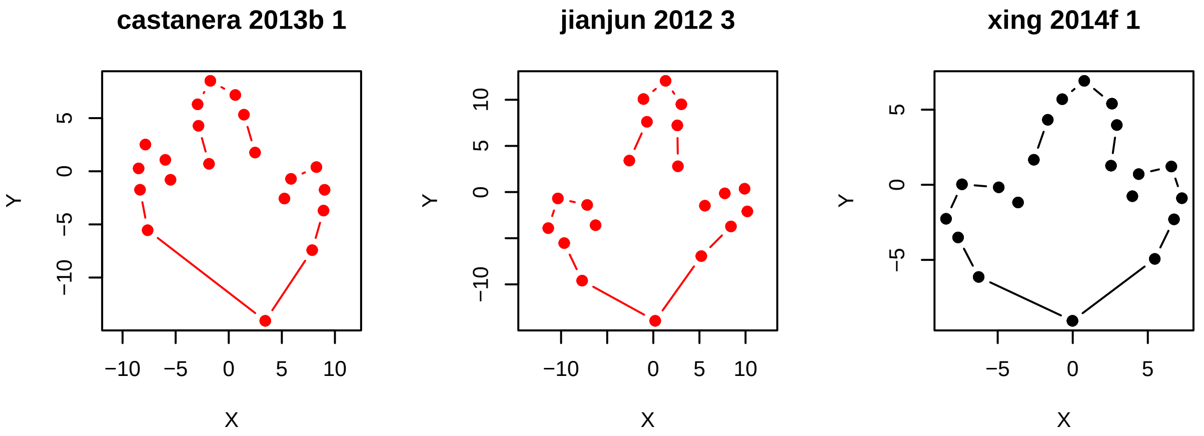


```
plotFootprint <- function(footprint, ...) {
    par(pty="s")

    plot(footprint,pch = 19, asp =1, ...)

    lmArc1 <- c(4, 7, 6, 5, 1, 8, 9)
    lmArc2 <- c(20, 19, 18, 2, 15, 16, 17)
    lmArc3 <- c(14, 13, 3, 10, 11, 12,4)

    lines(footprint[lmArc1, ], type="b", ...)

    lines(footprint[lmArc2, ], type="b",...)

    lines(footprint[lmArc3, ], type="b",...)
}
plotFootprint(footprint,
    col=ifelse(misclassIndicesGroup[i,"Group"]=="Theropoda", "black", "red"),
    main=rownames(misclassIndicesGroup)[i])
```

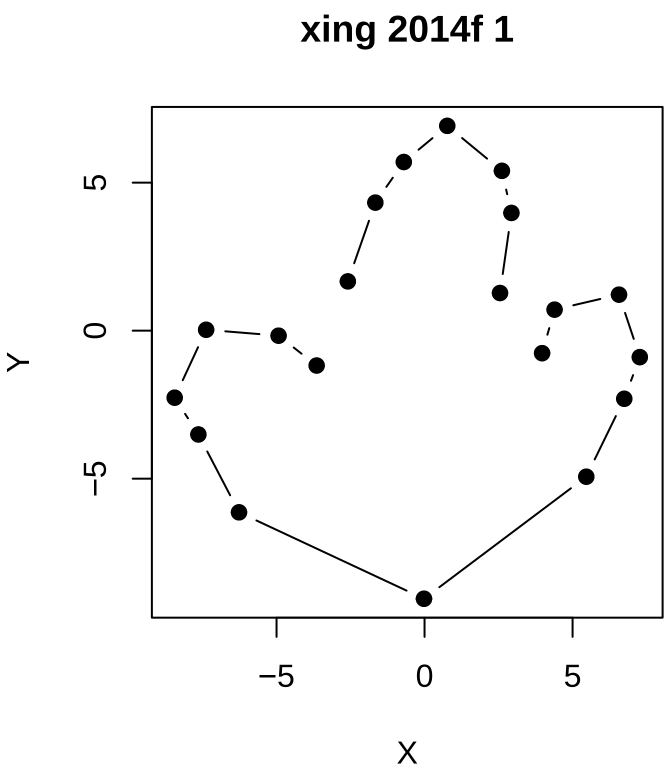


**Misclassified by 6 (= all) methods**

```
#, fig.width=14}
plotSomeFootprints <- function(dinoprints, group, indices, names, nCols=3, ...) {

  nTracks <- length(group)

  lenTrackVector <- ceiling(nTracks/nCols)

  #print(lenTrackVector)

  trackVector <- integer(lenTrackVector*nCols)
#print(trackVector)
  trackVector[1:nTracks] <- 1:nTracks

  # misclassIndicesGroup <- subset(misclassTable, subset = nMisclass==6, select = c(Group, indices))
  #print(trackVector)

layout(
  matrix(
    trackVector, ncol = nCols, byrow = TRUE)
)
#par(pty="s")
for (i in 1:nTracks ) {
  footprint <- subset(
      dinoprints, select = c(X, Y),
      subset = id==indices[i]
      )
  plotFootprint(footprint,
```

```
    col=ifelse(group[i]=="Theropoda", "black", "red")
    , main=names[i], ...
    )
}

}
plotSomeFootprints(dinoprints = dinoprints,
                   group = misclassTable[misclassTable$nMisclass==6, "Group"],
                   indices = misclassTable[misclassTable$nMisclass==6, "indices"],
                   names = rownames(misclassTable[misclassTable$nMisclass==6, ])
                   #, cex=3, cex.lab=2, cex.axis=2, cex.main=2
                   #, cex=2, cex.lab=2, cex.axis=2,  cex.main=2
                   )
```

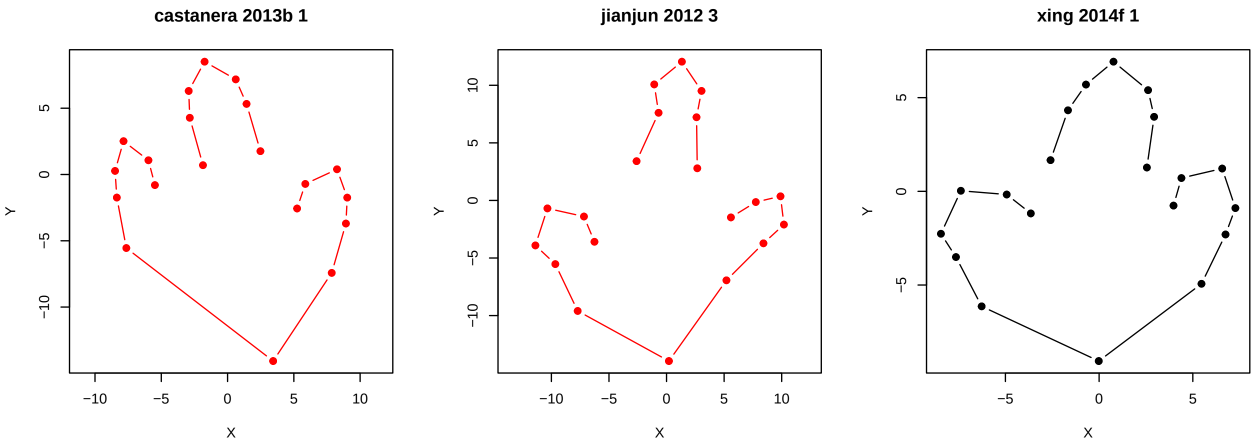


#### Misclassified by 5 methods

```
# misclassTable[misclassTable$nMisclass==5,]
plotSomeFootprints(dinoprints = dinoprints,
                   group = misclassTable[misclassTable$nMisclass==5, "Group"],
                   indices = misclassTable[misclassTable$nMisclass==5, "indices"],
                   names = rownames(misclassTable[misclassTable$nMisclass==5, ])
                   #, cex=3, cex.lab=2, cex.axis=2, cex.main=2
                   #, cex=2, cex.lab=2, cex.axis=2, cex.main=2
                   )
```

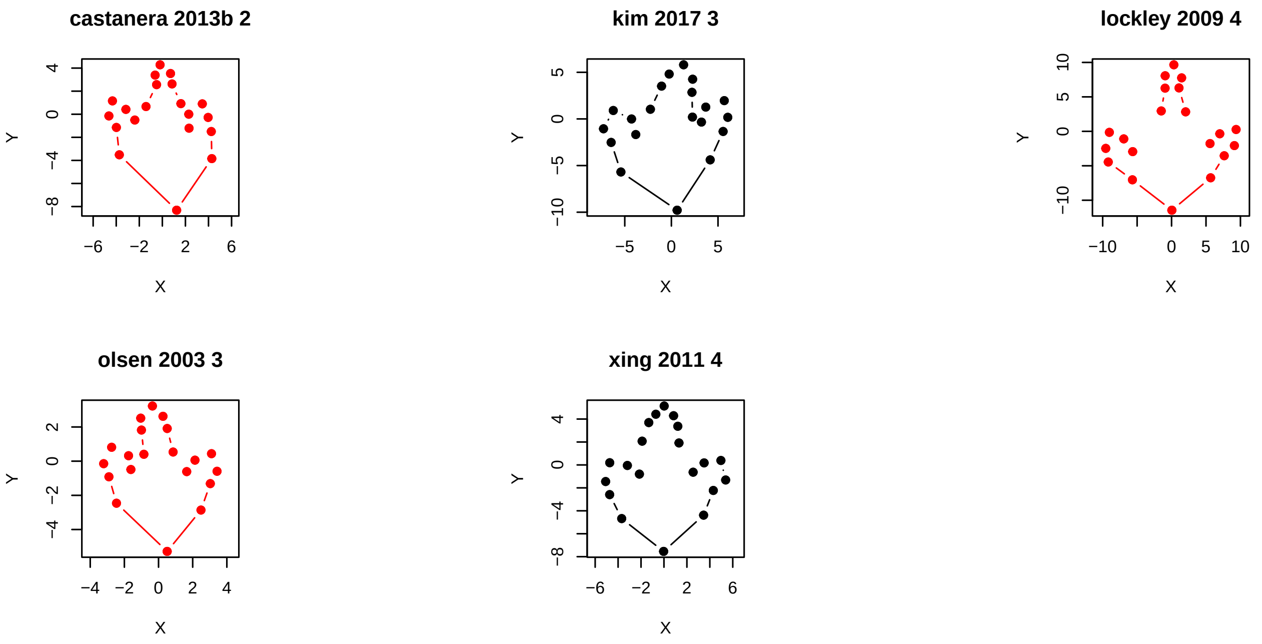


#### Misclassified by 4 methods

```r
#misclassTable[misclassTable$nMisclass==4,]
plotSomeFootprints(dinoprints = dinoprints, group = misclassTable[misclassTable$nMisclass==4, "Group"],
                   #,cex=2, cex.lab=2, cex.axis=2, cex.main=2
                   )
```

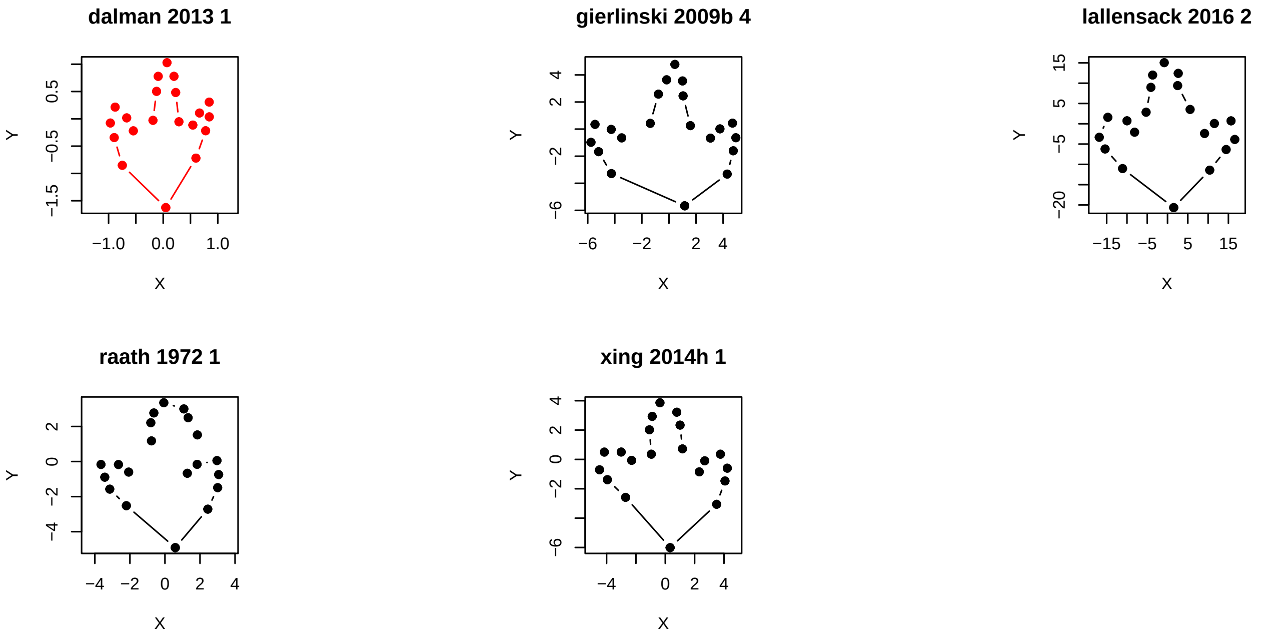


```r
# Plot misclassified samples to PDF
if(plotPDF) {
  basePrintNames <- gsub(" ", "_", rownames(misclassTable))
  for ( i in 1:nrow(misclassTable)) {
    printFN <- sprintf("figures/%s.pdf", basePrintNames[i] )
    footprint <- subset(
```

```
                   dinoprints, select = c(X, Y),
                   subset = id==misclassTable[i,"indices"]
                  )

    pdf(printFN)
    par(mar=c(4,4,1,1), cex=2)
    plotFootprint(footprint,
                  col=ifelse(
                    misclassTable[i,"Group"]=="Theropoda",
                    "black", "red"))
    dev.off()
  }
}
```
